# Single-cell transcriptomics reveals heterogenous thymic dendritic cell subsets with distinct functions and requirements for thymocyte-regulated crosstalk

**DOI:** 10.1101/2023.12.18.572281

**Authors:** Jayashree Srinivasan, Colin R. Moore, Aparna Calindi, Bryan R. Helm, Yilin Yang, John F. Moore, Hilary J Selden, Cody N. Heiser, Qi Liu, Ken S. Lau, Lauren I. R. Ehrlich

## Abstract

Dendritic cells are essential for establishing thymic central tolerance; however, mechanisms supporting their homeostasis and activation remain unresolved. Through single-cell transcriptomics and functional assays, we identify seven thymic conventional dendritic cell (cDC) subsets and discriminate their abilities to present self-antigens and induce regulatory T cells. Mice blocked at different stages of T-cell development reveal that CD4^+^ single-positive (SP) and CD8SP thymocytes differentially support homeostasis and activation of cDC1s versus cDC2s/plasmacytoid DCs (pDCs), respectively. CD8SPs indirectly support pDC survival and cDC2 thymic immigration, and they induce interferon signaling in cDCs, partly by promoting Type III interferon expression by medullary thymic epithelial cells (mTECs). In contrast, CD4SPs undergo cognate interactions with cDCs, inducing CD40 signaling required for activation of cDC1s. Activated cDC1s make non-redundant contributions to central tolerance. Altogether, this study comprehensively identifies distinct thymic DC subsets and elucidates requirements for crosstalk with thymocyte subsets that support their homeostasis, activation, and function.

## INTRODUCTION

Thymic central tolerance protects against autoimmunity by enforcing selection of self-tolerant T cells. If developing T cells express T-cell receptors (TCRs) with high affinity for self-antigens displayed by thymic antigen presenting cells (APCs), including medullary thymic epithelial cells (mTECs) and dendritic cells (DCs), they undergo negative selection or are diverted to the regulatory T cell (Treg) lineage, establishing central tolerance^1, 2^. Thymic DCs are crucial for central tolerance, as their genetic ablation results in impaired negative selection and autoimmunity^3^. In addition to displaying self-peptides from their proteome, DCs acquire and present diverse self-antigens to establish self-tolerance, including tissue-restricted antigens (TRAs) expressed by mTECs, circulating antigens, and peripherally-acquired antigens^4^.

The thymic DC compartment can be divided into three major cell types, cDC1s, cDC2s, and plasmacytoid DCs (pDCs)^5, 6^, with cDCs comprising non-activated CCR7^-^ MHC-II^lo^ and activated CCR7^+^MHC-II^hi^ subsets^7–11^. Thymic cDC2s are heterogeneous, underscoring the need for comprehensive assessment of thymic cDC composition and function^11–14^. Notably, cDC1s and cDC2s become activated in the sterile thymus environment, but our understanding of microenvironmental cues supporting their homeostasis and activation remains limited. While cDC1s differentiate from progenitors intrathymically, cDC2s and pDCs are recruited to the thymus^13–17^. Activation of DCs in the thymus mirrors peripheral DC activation, resulting in upregulation of MHC and co-stimulatory molecules that enhance self-antigen presentation to thymocytes^5, 8, 9^. Although critical for activation of peripheral DCs, pattern recognition receptor signaling is dispensable for thymic DC activation^8, 9^. Cognate interactions with CD4SP thymocytes are reported to support cDC1 and cDC2 activation, in part via CD40 signaling^9, 18^. Also, Type I and Type III interferons (IFNs), expressed by mTEC^hi^ cells, are needed to activate thymic cDC1s^19^, and type 2 cytokines support cDC2 homeostasis^11^. We previously showed that CCR7 expression supports survival of activated cDC1s (aDC1s)^7^. Multiple signals are required to fully license DCs in the periphery, raising the question of whether additional cellular or molecular cues regulate activation of thymic DC subsets.

Here, we employ single-cell RNA sequencing (scRNA-seq) and functional assays to identify transcriptionally distinct DC subsets in the mouse thymus, compare their APC activities, and discriminate mechanisms governing their homeostasis and activation. We find that aDC1s most efficiently present mTEC-derived antigens, followed closely by aDC2s. In contrast, cDC2s and aDC2s most efficiently induce Treg differentiation *in vitro*. Notably, CD4SPs support homeostasis of cDCs and activation of cDC1s, while CD8SPs are required for homeostasis of pDCs and cDC2s, as well as activation of cDC2s. CD4SPs undergo cognate interactions with cDC1s, driving CD40 signaling that is critical for cDC1 activation. In contrast, CD8SPs alter the thymus environment, promoting IFN expression by mTEC^hi^ cells and activating cDCs via IFN signaling. CD8SPs also indirectly support intrathymic pDC survival and recruitment of cDC2s to the thymus. CD40-activated aDC1s play a non-redundant role in supporting early-stage negative selection and Treg induction. Altogether, our study demonstrates substantial phenotypic and functional heterogeneity in the thymic DC compartment, identifies differential thymocyte-mediated signals that support homeostasis and activation of DC subsets, and demonstrates the impact of aDC1s on central tolerance.

## RESULTS

### scRNA-seq reveals transcriptionally distinct thymic cDC subsets, which localize to different thymic regions

To identify transcriptionally distinct thymic DC subsets, we performed scRNA-seq analysis of 8404 FACS-purified CD45^+^MHC-II^+^ thymic hematopoietic APCs (HAPCs) from 1 month-old C57BL/6J mice. Distinct HAPC subsets and thymocytes were identified by expression of lineage-specific genes (Extended Data Fig. 1a-c). cDCs were re-clustered and analyzed separately, enabling identification of 11 transcriptionally distinct cDC subsets: 5 cDC1 clusters expressing *Xcr1, Irf8,* and *Cd36*^11, 20, 21^, 3 cDC2 clusters expressing *Sirpa*, *Irf4,* and *Mgl2*^11, 20^, a heterogenous “Ly6d”^+^ cDC cluster expressing *Zbtb46, Xcr1,* and *Sirpa,* and 2 activated “aDC” clusters expressing DC maturation genes, such as *Ccr7*, *Cd63*, *Ccl5,* and *Ccl22* (Fig. 1a,b and Extended Data Fig. 1d,e)^8, 9^.

**Figure 1.**
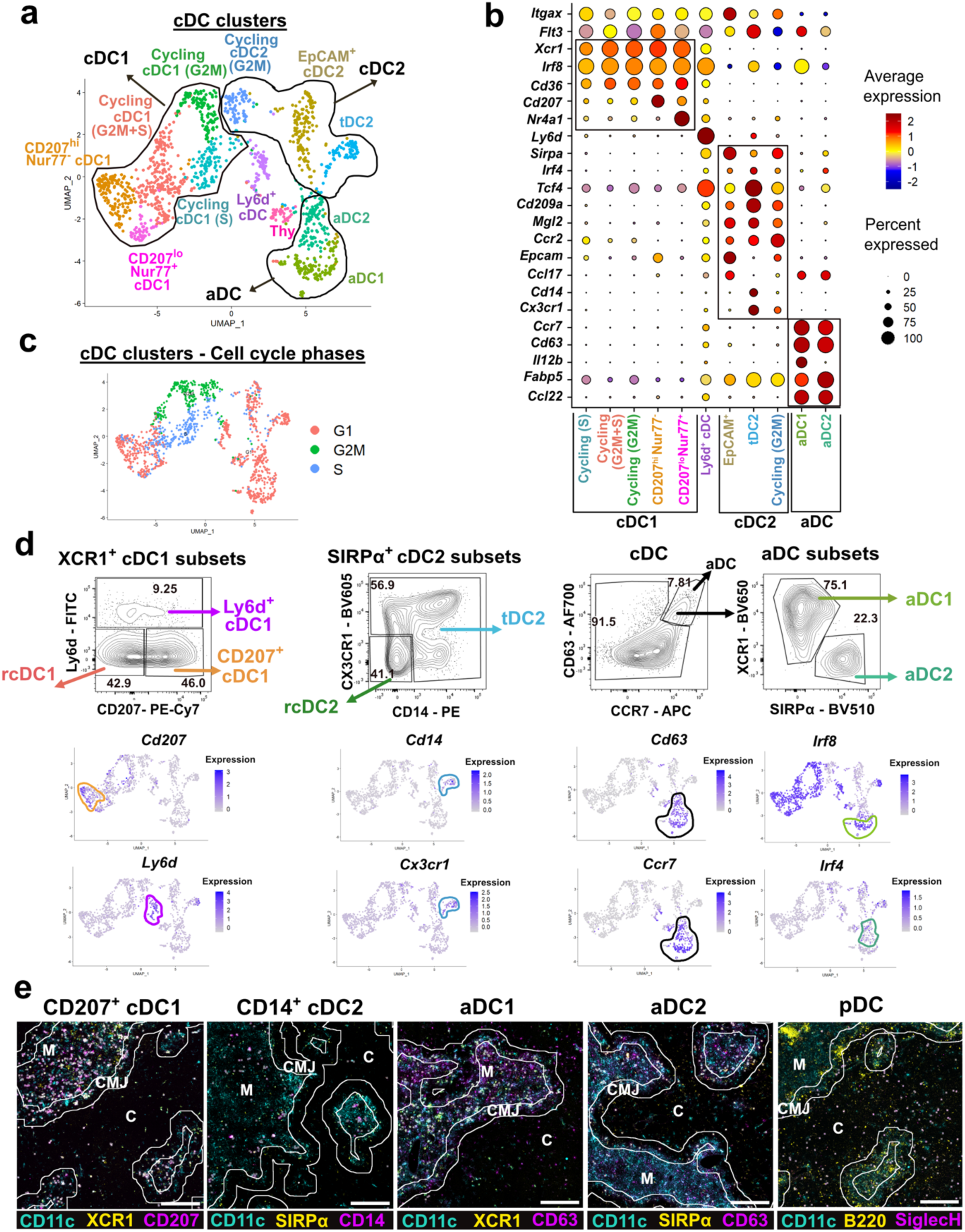
scRNA-seq reveals transcriptional heterogeneity in the thymic DC compartment. (a) UMAP visualization of 1399 re-clustered thymic cDCs from adult C57BL/6J (WT) mice (n=2) depicting 11 transcriptionally distinct cDC clusters. (b) Normalized expression (z-score) of select DC marker genes across the transcriptionally defined cDC clusters from (a). The transcriptional clusters have been ordered as *Ccr7^-^*cDC1 clusters, *Ccr7^-^* cDC2 clusters, and *Ccr7^+^* activated “aDC” clusters. (c) Cell-cycle phases across the cDC clusters from (a). (d) (Top) Representative flow cytometry plots identifying 7 distinct cDC subsets based on protein expression of cluster-specific genes (gating strategy in Extended Data Fig. 2a). (Bottom) Expression levels of cDC cluster-specific genes for the respective flow-defined subsets above. (e) Representative immunofluorescence used to quantify locations of the indicated DC subsets in thymic sections (representative of N=3 independent experiments). Scale bar, 250µm. Lines demarcate the cortex (C), medulla (M), and CMJ in each thymic section. Centroids were overlaid on cells meeting the criteria for each indicated cDC subset (see Methods). See also Extended Data Fig. 1 and 2.

cDC1 clusters comprise “CD207^hi^ Nur77^-^ cDC1s” expressing *Cd207* and *Itgae*, three “cycling cDC1” clusters expressing genes associated with cell proliferation, and “CD207^lo^ Nur77^+^ cDC1s” expressing high levels of *Nr4a1* and AP-1 family genes^22^, including *Jun* and *Fos* (Fig. 1b,c and Extended Data Fig. 1d). cDC2 clusters comprise “cycling cDC2s”, “EpCAM^+^ cDC2s” expressing *Epcam* and *Ccl17,* and a recently described transitional “tDC2” subset^23, 24^ expressing pDC (e.g, *Tcf4, Siglech)* and monocyte-associated (e.g.*Cx3cr1, Cd14*) genes (Fig. 1b and Extended Data Fig. 1d), reminiscent of cDC2 subsets that traffic peripheral self-antigens into the thymus or capture blood-borne antigens to induce tolerance^12–14^. Lastly, while sharing a strong DC activation signature, lineage-specific genes discriminate aDC1s (e.g., *Irf8, Il12b*) from aDC2s (e.g., *Irf4, Fabp5*), correlating with previously reported CCR7^+^ mature “mDC1” and “mDC2” clusters, respsectively^11^ (Fig. 1b and Extended Data Fig. 1d,f).

We established a flow cytometry panel to distinguish thymic DC subsets identified by scRNA-seq (Extended Data Fig. 2a). We identified cDCs as F4/80^-^PDCA1^-^ CD11c^+^MHC-II^+^ cells within the myeloid (CD11c^+^ and/or CD11b^+^) compartment. Activated cDCs co-express CCR7 and CD63, while XCR1 versus SIRPα expression delineates aDC1 versus aDC2 subsets, respectively (Fig. 1d). Non-activated CCR7^-^CD63^-^ cDCs are divided into cDC1 and cDC2 subsets by expression of XCR1 versus SIRPα, respectively (Extended Data Fig. 2a). Three cDC1 subsets were identified by flow cytometry: CD207^+^ cDC1s, Ly6d^+^ cDC1s, and CD207^-^Ly6d^-^ cDC1s, annotated as “remaining” cDC1s (rcDC1) (Fig. 1d), which comprise cycling and CD207^lo^Nur77^+^ cDC1 transcriptional clusters (Fig. 1d). Two cDC2 subsets were identified by flow cytometry: CD14^+^ and/or CX3CR1^+^ tDC2s and CD14^-^CX3CR1^-^cDC2s, denoted “remaining” cDC2s (rcDC2), comprising cycling and EpCAM^+^ cDC2 transcriptional clusters (Fig. 1d).

To validate that the 7 cDC subsets identified by flow cytometry correlate with their transcriptional counterparts, cDC subsets were FACS-isolated, hashtagged, and pooled prior to scRNA-seq analysis. The 6744 hashtagged cDCs were distributed among 10 transcriptionally distinct clusters (Extended Data Fig 2b,c). Six FACS-isolated cDC subsets mapped to expected cDC transcriptional cluster(s) (Extended Data Fig. 2d,e), but insufficient numbers of Ly6d^+^ cDC1s prevented accurate annotation of this subset. Sorted CCR7^+^ CD63^+^ aDC1s and aDC2s mapped almost exclusively to the expected aDC clusters (Extended Data Fig. 2e).

Because thymocytes migrate between distinct microenvironments, where local APCs impact their differentiation and selection^25^, we assessed the localization of thymic DC subsets (Fig. 1e and Extended Data Fig. 2f,g). Overall, cDCs and aDCs are most abundant in the medulla followed by the cortico-medullary junction (CMJ). CD14^+^ tDC2s are located almost exclusively in the medulla (Fig. 1e and Extended Data Fig. 2g,h). Consistent with localization of thymic macrophages^26^, 83.1% of cortically localized SIRPα^+^CD63^+^ cells expressed MertK; allowing us to correct for macrophages in aDC2 density calculations (Extended Data Fig. 2f,h). pDCs are the most abundant DC subset in the cortex, followed by CD207^+^ cDC1s, which may thus interact with early post-positive selection thymocytes^27–29^ (Fig. 1e and Extended Data Fig. 2g-i). Altogether, thymic DC subsets localize in distinct anatomical niches, where they could interact with co-localized thymocyte subsets to establish robust central tolerance.

### cDC subsets have distinct capacities to present mTEC-derived antigens and induce Treg differentiation

We assayed thymic DC subsets for distinct functional activities. Previous reports showed that aDC1s are best able to acquire antigens expressed by AIRE^+^ mTECs^10^, and they present mTEC-derived antigens on MHC-I and MHC-II more efficiently than non-activated cDC1s^8^. However, the relative capacities of the delineated cDC subsets identified herein to acquire and present mTEC-derived antigens on MHC-I and MHC-II remains unclear. To test for cross-presentation of mTEC-derived TRAs on MHC-I, we co-cultured OT-I CD8^+^ T cells with thymic DC subsets FACS-sorted from RIP-mOVA and RIP-OVA^hi^ transgenic mice^30, 31^, in which mTECs express ovalbumin (OVA) as a model TRA. With both membrane-bound (RIP-mOVA) and secreted (RIP-OVA^hi^) TRA models, aDC1s most efficiently cross-presented the TRAs on MHC-I, inducing proliferation of OT-I CD8^+^ T-cells nearly equivalent to splenic APCs presenting exogenously supplied OVA peptide (Fig. 2a-c and Extended Data Fig. 3a-c). Although inferior to aDC1s (p-value <0.0001), aDC2s cross-presented OVA to stimulate more OT-I proliferation than non-activated cDC1 or cDC2 subsets (Fig. 2a-c and Extended Data Fig. 3a-c).

**Figure 2.**
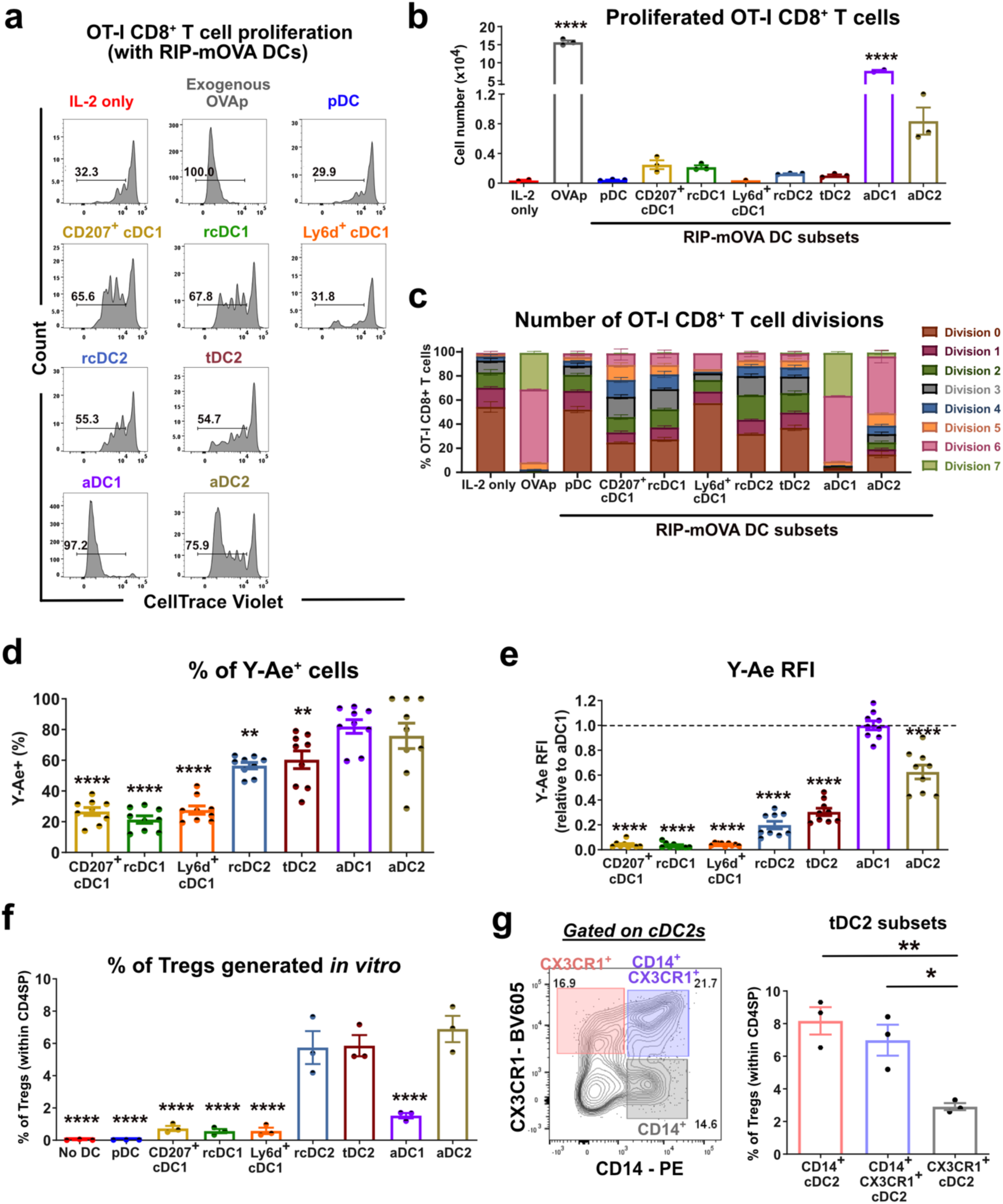
cDC subsets have distinct abilities to present antigens and induce Treg differentiation. (a) Representative histograms, with gating shown to identify the percentage of OT-I CD8^+^ T cells that underwent at least 1 cell division after co-culture for 3.5 days with the specified DC subsets from RIP-mOVA mice. OT-I CD8^+^ T cells incubated only with IL-2 served as negative controls, while those co-cultured with splenocytes pulsed with 50nM of SIINFEKLp (OVAp_257-264_) served as positive controls. For (a-c), data are representative of 3 independent experiments. (b) Quantification of OT-I CD8^+^ T cells that underwent proliferation from the experiment in (a). Bars represent mean ± SEM. Each dot represents a replicate well. (c) Modeling, based on the experiment in (a), of the frequency of OT-I CD8^+^ T cells that underwent the indicated number of cell divisions. Bars represent mean ± SEM. (d) Frequency of Y-Ae^+^ cells and (e) relative fluorescence intensity (RFI) of Y-Ae expression on C57BL/6J donor-derived thymic cDC subsets from BM chimeras of C57BL/6J hematopoietic progenitors transplanted into lethally irradiated BALB/c mice. (d-e) Data are compiled from N= 3 independent experiments (n=9 mice total). Bars represent mean ± SEM and symbols represent individual mice. (e) Data are normalized to the average Y-Ae MFI of aDC1s in each individual experiment. (f) Percentage of Foxp3^+^ CD25^+^ Tregs induced after culture of CD73^-^ CD25^-^ Foxp3^-^ CD4SP thymocytes with the indicated C57BL/6J thymic DC subsets for 4 days. (f-g) Data are compiled from N= 3 independent experiments. Bars represent mean ± SEM and symbols represent average of triplicate wells per DC subset from each experiment. (g) (Left) Representative flow cytometry plot depicting the three heterogenous subpopulations within tDC2s. (Right) Percentage of Tregs induced by each tDC2 subpopulation in an in vitro Treg generation assay, as in (f). Statistical analysis was performed using one-way ANOVA with (b, d, e, and f) Dunnett’s multiple comparisons test and (g) Tukey’s multiple comparisons test; *p<0.05, ** p<0.01, ****p<0.0001. Significance is relative to (b) “IL-2 only” control, (d and e) aDC1 subset, and (f) aDC2 subset. See also Extended Data Fig. 3.

Next, we compared the ability of cDC subsets to present mTEC-derived antigens on MHC-II. C57BL/6J bone marrow (BM)-derived hematopoietic progenitors were transplanted into lethally irradiated BALB/c recipient mice, yielding chimeric thymuses in which only radio-resistant BALB/c cells express I-Eα, while only hematopoietic cells express I-A^b^. Thus, cDCs must acquire I-Eα from mTECs for presentation on I-A^b^, and the Eα:I-A^b^ complex can be detected by the Y-Ae antibody^8, 32^. A high proportion of both aDC1s and aDC2s presented mTEC-derived Eα on MHC-II, with aDC1s displaying the highest levels of Eα:I-A^b^ complexes, followed by aDC2s and then rcDC2s and tDC2s (Fig. 2d,e).

We next tested the capacity of different DC subsets to induce Treg differentiation. cDC subsets were FACS-purified from C57BL/6J thymi and co-cultured with sorted CD73^-^ CD25^-^Foxp3^-^ newly generated CD4SP thymocytes^33, 34^. Although aDC1s present the highest levels of mTEC-derived antigens on MHC-II (Fig. 2e), rcDC2s, tDC2s, and aDC2s were significantly more efficient at inducing Tregs relative to other DC subsets (Fig. 2f). Furthermore, CD14^+^ tDC2s generated Tregs more efficiently than CD14^-^ tDC2s (Fig. 2g). We considered whether differential expression of genes associated with antigen processing and presentation might underlie functional differences between cDC subsets. aDC1s express high levels of genes associated with MHC-I antigen processing and presentation (*Psme1, Tapbp*, *Tap1*, *Tap2*), possibly contributing to their superior ability to present antigens on MHC-I. In contrast, aDC2s express higher levels of genes associated with MHC-II antigen processing and presentation (*Cd74, Ciita, Ctsh, H2-Oa,* and *H2-DMa* (Extended Data Fig. 3d)), possibly increasing the diversity of self-antigens presented and enabling continued expression of new MHC-II molecules. Consistent with DC activation in the periphery^35^, thymic aDC1s and aDC2s express high cell-surface levels of MHC-I, MHC-II, and the costimulatory molecules CD80 and CD86 (Extended Data Fig. 3e,f,g), which are indispensable for thymocyte negative selection and Treg induction^36^.

Altogether, while aDC1s, closely followed by aDC2s, most efficiently present mTEC-derived antigens on MHC-I and MHC-II, cDC2s and aDC2s more efficiently induce Treg generation in vitro.

### Thymic cDC1 and cDC2 homeostasis and activation depend on the presence of distinct thymocyte subsets

It is established that cDC1s differentiate from progenitors intrathymically, while cDC2s migrate into the thymus^5, 6^, but the cellular and molecular cues required for homeostasis and activation of DC subsets within the thymus remain unresolved. Presentation of self-antigens to CD4SP thymocytes has been implicated in activation of cDC1s and cDC2s^9, 18^. Given the significant heterogeneity of thymic DCs revealed by scRNA-seq, we sought to determine if different thymocyte cues regulate homeostasis and activation of distinct cDC subsets.

We first analyzed the thymic cDC compartment in *Rag2^-/-^* and *Tcra^-/-^* mice, in which thymocyte differentiation is blocked at the CD4^-^CD8^-^CD25^+^CD44^-^ double-negative 3 (DN3)^37^ and pre-positive selection CD4^+^CD8^+^ double-positive (DP) stages^38^, respectively. In both strains, the cellularity of all cDC subsets and pDCs declined (Fig. 3a,b), with a substantial drop in aDC1s and aDC2s (Fig. 3a,b and Extended Data Fig.4a,b), indicating that post-positive selection thymocytes are required for homeostasis and activation of both cDC1s and cDC2s.

**Figure 3.**
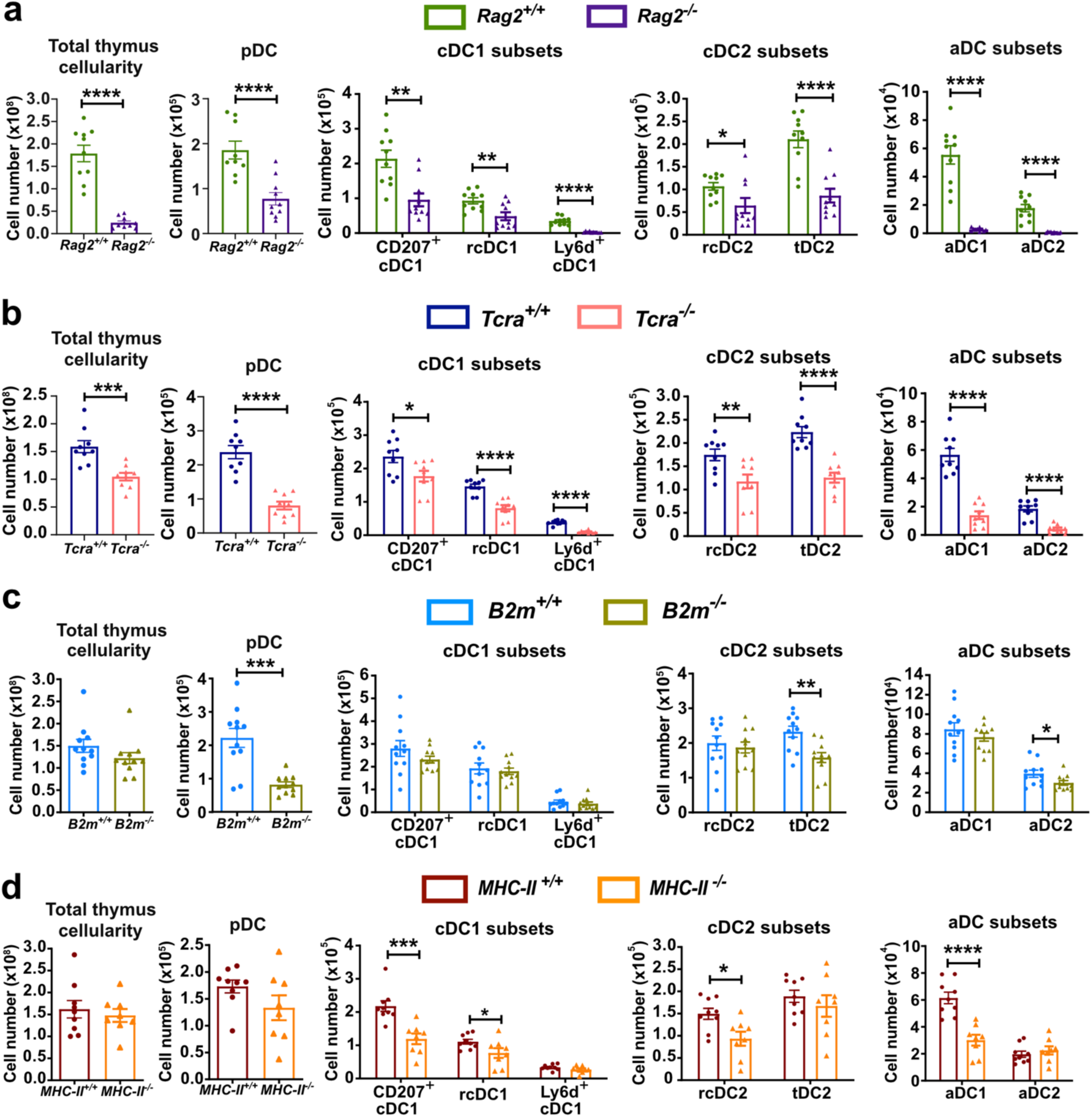
Thymic cDC1 and cDC2 homeostasis and activation depend on the presence of distinct thymocyte subsets. (a-d) Quantification of (left to right) total thymus cellularity and cell numbers of pDCs, cDC1 subsets, cDC2 subsets, and aDC subsets from the thymi of WT versus (a) *Rag2^-/-^*, (b) *Tcra^-/-^*, (c) *B2m^-/-^* and (d) *MHC-II^-/-^* littermate mice. (a-d) Data are compiled from N=3-4 independent experiments (n=8-11 mice). Bars represent mean ± SEM and symbols represent individual mice. Statistical analysis was performed using unpaired Student *t*-test; *p<0.05, ** p<0.01, ***p<0.001, ****p<0.0001. See also Extended Data Fig. 4 and Extended Data Fig. 5.

*B2m^-/-^* mice were analyzed to evaluate the contribution of CD8SP thymocytes to pDC and cDC homeostasis and activation. β2m-deficiency resulted in fewer pDCs and tDC2s (Fig. 3c and Extended Data Fig. 4c), highlighting a role for CD8SP thymocytes in their homeostasis. Interestingly, the number of aDC2s also declined significantly in *B2m^-/-^*mice (Fig. 3c). Since *B2m^-/-^* mice lack both invariant natural killer T cells (iNKT) and CD8SP thymocytes^39^, we tested if iNKT deficiency could account for the altered DC compartment in *Β2m^-/-^* mice, but found no deviation in thymic DC numbers in *Cd1d^-/-^* mice (Extended Data Fig. 4d). These results indicate that CD8SP thymocytes support pDC and tDC2 homeostasis and cDC2 activation.

Thymic DCs in *MHC-II^-/-^* mice were analyzed to test the contribution of CD4SP thymocytes to DC homeostasis and activation. pDC numbers remained unaltered in the absence of CD4SPs, confirming a non-redundant role for CD8SPs in regulating pDC homeostasis (Fig. 3d). CD207^+^ cDC1s and rcDC1s declined significantly in the absence of CD4SP thymocytes, as did rcDC2s (Fig. 3d and Extended Data Fig. 4e). Contrary to previous findings^9^, while CD4SP deficiency resulted in a significant decline in aDC1s, aDC2s were intact (Fig. 3d and Extended Data Fig. 4e). Of note, the decline in aDCs was more pronounced in *Rag2*^-/-^ and *Tcra*^-/-^ mice than in mice deficient for either individual SP subset (Fig. 3a-d), indicating some redundancy in the ability of CD4SP and CD8SP thymocytes to support thymic DC activation.

To determine if thymic DC functionality is impaired in the absence of thymocyte subsets, we tested if cDCs from *Rag2^-/-^*, *B2m^-/-^*, or *MHC-II^-/-^* thymi could induce Treg differentiation in vitro. cDC subsets from *Rag2^-/-^* and *B2m^-/-^*thymi induced Tregs comparably to their WT counterparts (Extended Data Fig. 4f-g). Consistent with a previous study^34^, cDC subsets from *MHC-II^-/-^* mice were unable to induce Treg differentiation (Extended Data Fig. 4h), demonstrating that antigen presentation on MHC-II is required for in vitro Treg induction. Together, these results show that not all DC activity involved in central tolerance requires thymocyte crosstalk.

Because pDCs and cDC2s migrate into the thymus from the periphery^14–16, 40^, we tested whether the decline in these subsets in the absence of thymocyte crosstalk could reflect altered DCs in secondary lymphoid organs (SLOs). The frequency of pDCs declined in SLOs of *Rag2^-/-^* mice but remained unaltered in *B2m^-/-^* mice (Extended Data Fig. 5a-d), indicating that the reduction in thymic pDCs in *B2m^-/-^* mice (Fig. 3c) likely reflects intrathymic changes in pDC homeostasis. Moreover, the frequency of tDC2s increases in LNs of *B2m^-/-^* mice (Extended data Fig. 5c); thus, the reduction in thymic tDC2s does not reflect a peripheral decline in this subset. The frequency of cDC2s was reduced in *MHC-II^-/-^* SLOs (Extended Data Fig. 5e,f), but not in the thymus of these mice (Extended Data Fig. 4e), indicating that CD4^+^ T cells are required for peripheral cDC2 homeostasis.

Altogether, our findings show that CD4SP thymocytes primarily regulate cDC1 homeostasis and activation while CD8SP thymocytes regulate homeostasis of pDCs, tDC2s, and aDC2s.

### cDC1 homeostasis and activation require cognate interactions with CD4SP thymocytes, while CD8SP thymocytes indirectly regulate pDC and tDC2 homeostasis

CD4SP and CD8SP thymocytes may modulate cDC homeostasis and activation via direct interactions with DC subsets or via indirect effects on the thymus microenvironment. Notably, SP thymocytes are indispensable for maturation of AIRE^+^ mTECs^41, 42^, which produce cytokines that regulate DC activation^19, 43^.

To distinguish between direct and indirect CD4SP -mediated regulation of DCs, we generated mixed BM chimeras in which a mixture of congenically distinct WT and *MHC-II^-/-^* hematopoietic progenitors were transplanted into lethally irradiated WT mice (Fig. 4a,b). In host thymi, CD4SP thymocyte differentiation was intact and MHC-II-expressing WT-derived HAPCs were present, contributing to a normal thymus environment. After 10 weeks, thymocyte chimerism from WT versus *MHC-II^-/-^* donor cells was comparable (Fig. 4a). However, *MHC-II^-/-^* DCs of all subsets, except pDCs and aDC2s, were underrepresented (Fig. 4b). Thus, when competing with WT DCs, MHC-II expression is required in a cell-intrinsic manner for cDC1 and cDC2 homeostasis. Consistent with global *MHC-II^-/-^*mice (Fig. 3d; Extended Data Fig. 4e), *MHC-II*^-/-^ cells contributed poorly to aDC1s, but efficiently generated aDC2s (Fig. 4b). These data suggest that cDC1s and cDC2s require MHC-II-mediated interactions with CD4SPs for their homeostasis, but only cDC1s require interactions with CD4SPs for activation.

**Figure 4.**
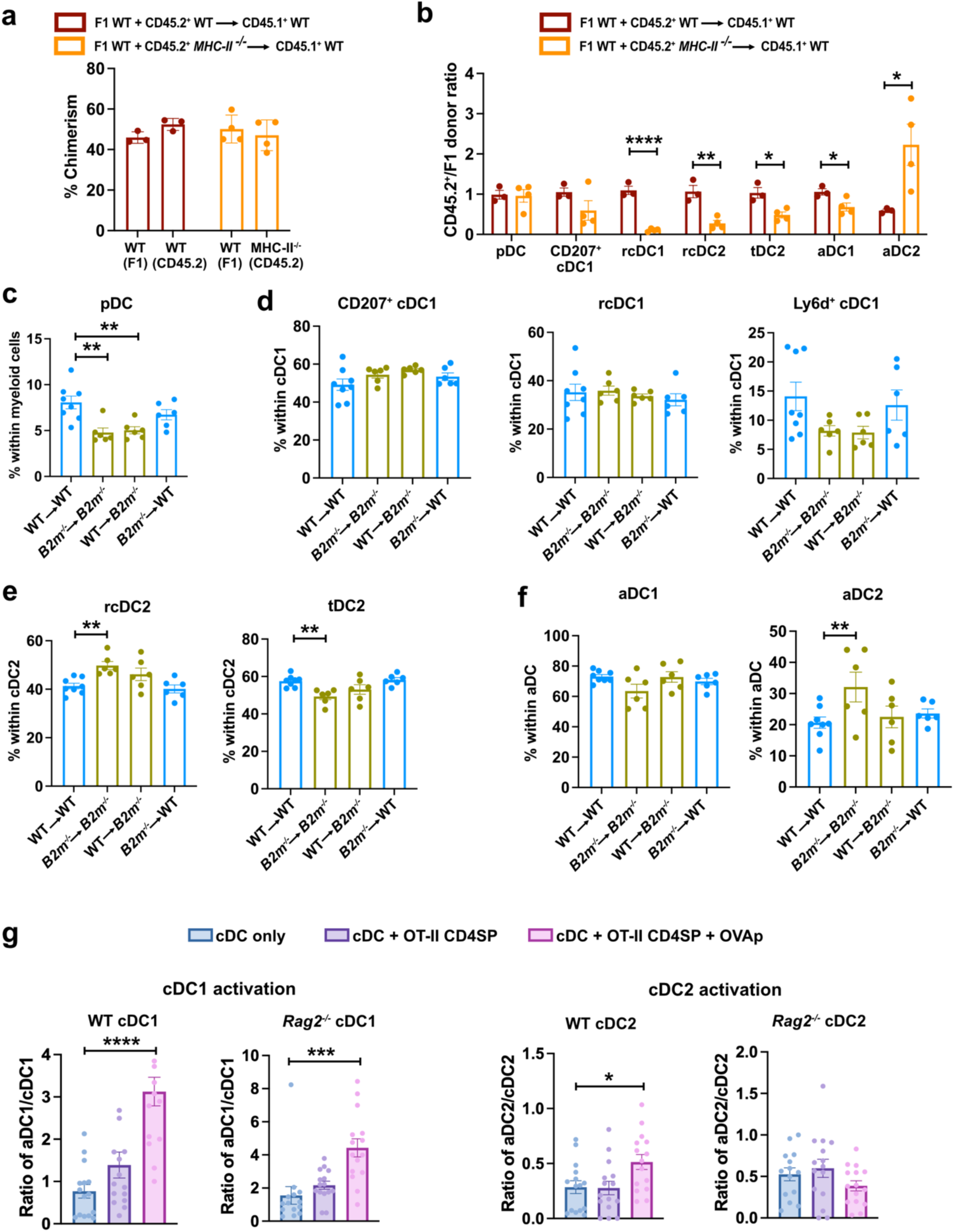
cDC1 homeostasis and activation requires cell-intrinsic MHC-II expression, implicating direct interactions with CD4SPs, while pDC homeostasis is regulated indirectly by the presence of CD8SPs. (a) Overall donor thymic chimerism within mixed BM chimera recipients in which a 1:1 mixture of F1 WT and either CD45.2^+^ WT or CD45.2^+^ *MHC-II*^-/-^ BM were transferred into lethally irradiated CD45.1^+^ WT mice. Bars represent mean ± SEM. (b) Quantification of the ratios of CD45.2^+^/ F1 donor cells within each indicated thymic DC subset from mixed BM chimeras in (a). (a-b) Data are compiled from N=2 independent experiments (n=3-4 mice). (c) The frequency of pDCs within the thymic myeloid compartment of the indicated reciprocal BM chimera recipients in which WT or *B2m^-/^*^-^ hematopoietic progenitors were transplanted into lethally irradiated recipients of either genotype. (d-f) The frequency of (d) cDC1, (e) cDC2, and (f) aDC subsets within the indicated thymic DCs populations of reciprocal *B2m^-/-^* BM chimera recipients. (c-f) Data are compiled from N=3 independent experiments (n=6-8 mice). Bars represent mean ± SEM. (g) Ratio of activated to non-activated cDC1s and cDC2s identified 17-19 hours after non-activated FACS-purified PDCA1^-^F4/80^-^CD11c^+^CCR7^-^CD63^-^MHC-II^lo^ cDC1s or cDC2s from WT or *Rag2*^-/-^ thymi were cultured alone (blue), with OT-II CD4SPs (purple), or with OT-II CD4SPs in the presence of 1uM OVAp_323-339_ (pink). Data are compiled from N= 5 independent experiments. Bars represent mean ± SEM and each symbol represents data from one replicate well. Statistical analysis was performed using (a,b) unpaired Student *t*-test and (c-f) one-way ANOVA with Dunnett’s multiple comparisons corrections relative to WT into WT or (g) respective cDC only control; *p<0.05, ** p<0.01, ***p<0.001, ****p<0.0001.

To determine whether CD8SP thymocytes directly or indirectly support pDC/cDC2 homeostasis, we analyzed reciprocal BM chimeras in which WT or *B2m*^-/-^ hematopoietic progenitors were transplanted into recipients of either genotype (Fig. 4c-f). Consistent with constitutive *B2m^-/-^* mice (Extended Data Fig. 4c), thymic pDC and tDC2 subsets declined when both donor and host were *B2m*-deficient, while other DC subsets were unimpaired (Fig. 4c-f). Interestingly, pDCs declined when hematopoietic progenitors of either genotype were transplanted into *B2m*-deficient hosts, indicating pDC homeostasis requires MHC-I expression by radioresistant thymic stroma (Fig. 4c). A similar, albeit non-significant, trend was observed for tDC2s (Fig. 4e). Because positive selection of CD8SP thymocytes is impaired in *B2m*^-/-^ hosts, these data suggest an indirect role for CD8SP thymocytes in supporting pDC and tDC2 homeostasis, while cognate interactions between MHC-I-expressing DCs and CD8SPs are dispensable.

The absence of positively selected thymocytes in *Rag2^-/-^*and *Tcra*^-/-^ mice caused profound impairment in cDC1 and cDC2 activation (Fig. 3a,b). To test whether deprivation of SP thymocyte-derived crosstalk results in sustained cDC dysfunction, we established an in vitro thymic DC activation assay. CCR7^-^ CD63^-^ MHC-II^lo^ non-activated DCs were incubated with OT-II TCR transgenic CD4SP thymocytes^44^ in the presence or absence of cognate OVA peptide (Fig. 4g). Notably, WT cDC1s and cDC2s were activated only when presenting OVA peptide to OT-II CD4SP thymocytes. While cognate interactions with OT-II CD4SP thymocytes were sufficient to rescue *Rag2^-/-^* cDC1 activation, *Rag2^-/-^* cDC2s failed to become activated (Fig. 4g), indicating sustained impairment of cDC2s in the absence of thymocyte-derived signals (Fig. 4g).

Altogether, analyses of BM chimeras indicate that cognate interactions with CD4SPs are indispensable for cDC1 and cDC2 homeostasis and cDC1 activation, but are not required for cDC2 activation. In contrast, CD8SP thymocytes indirectly regulate thymic pDC and tDC2 homeostasis.

### scRNA-seq profiling identifies the presence of unique DC transcriptional states in the absence of thymocyte crosstalk

To identify candidate mechanisms underlying thymocyte support of DC homeostasis and activation, we performed scRNA-seq on HAPCs from *MHC-II^-/-^*, *B2m^-/-^* and *Rag2^-/-^* thymi (Extended data Fig. 6a,b). Re-clustering cDCs separately revealed 17 transcriptionally unique cDC subsets across genotypes: 7 cDC1, 6 cDC2, 1 Ly6d^+^ cDC, and 3 aDC subsets (Fig. 5a,b and Extended Data Fig. 6c), indicating greater heterogeneity than in WT cDCs (Fig. 1a). Expected genotype-specific gene expression changes in knockout (KO) cDCs were confirmed (Extended Data Fig. 6d,e).

**Figure 5.**
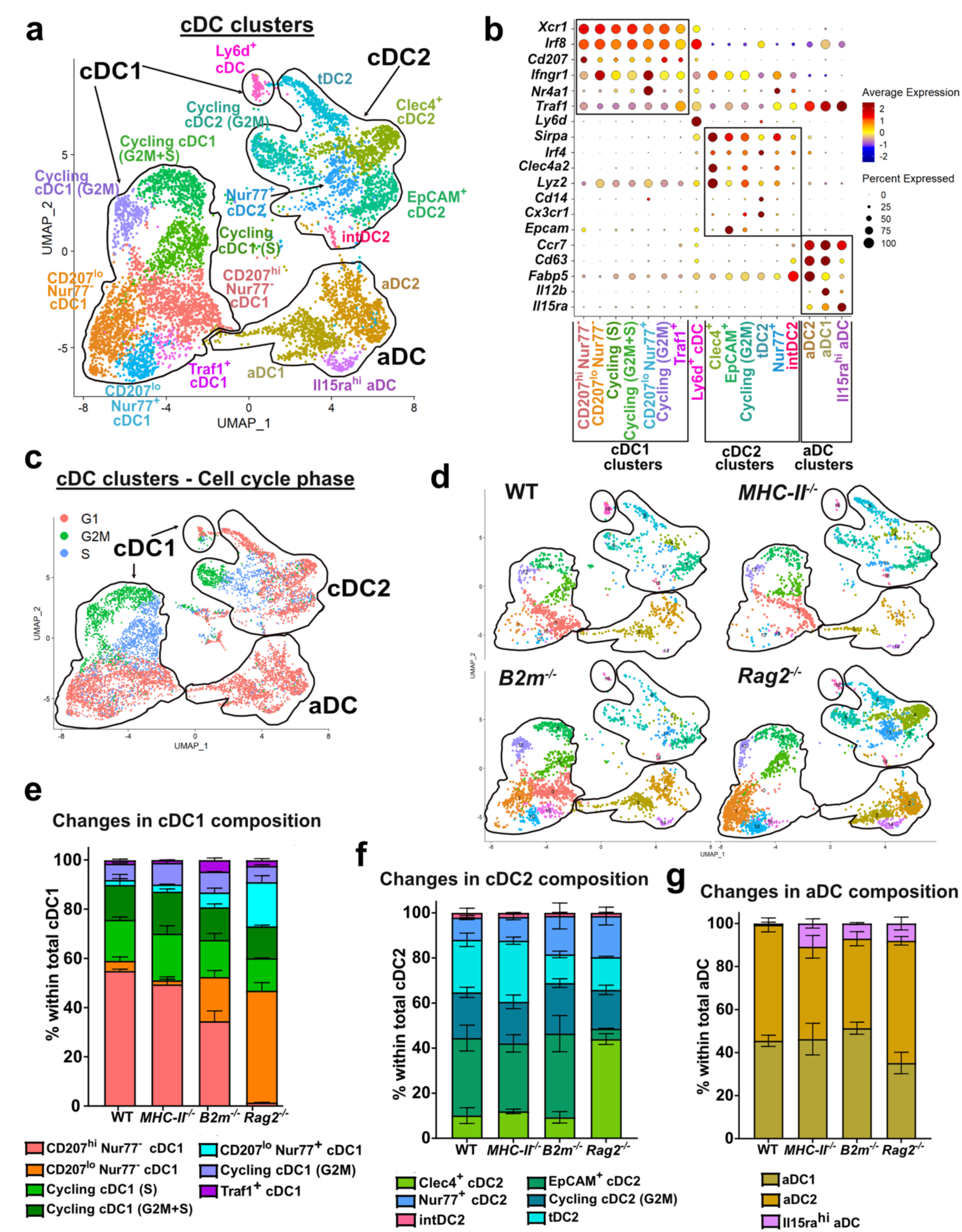
scRNA-seq profiling identifies the emergence of unique DC transcriptional states in the absence of thymocyte crosstalk. (a) Merged UMAP representation of 7982 re-clustered cDCs from WT, *MHC-II^-/-^*, *B2m^-/-^*and *Rag2^-/-^* thymi (n=2 mice per genotype). (b) Dot plots showing normalized expression (z-score) of cDC1-, cDC2- and aDC-specific genes across cDC clusters from (a). (c) Cell-cycle phase analysis for cDC clusters from (a). (d) UMAP representation of cDC clusters from (a), separated by genotype of origin. (e-g) Frequencies of transcriptionally distinct (e) cDC1 clusters, (f) cDC2 clusters, and (g) aDC clusters shown in (a), within the parent population indicated on the Y-axis label separated by the indicated genotypes (n=2 mice per genotype). Bars represent mean ± SEM. See also Extended Data Fig. 6.

Five cDC1 clusters matched those in WT thymi (Fig. 1a): CD207^hi^Nur77^-^ cDC1, 3 cycling cDC1 subsets, and CD207^lo^ Nur77^+^ cDC1 (Fig. 5b,c). Two cDC1 clusters were primarily detected in KO genotypes (Fig. 5d,e): “CD207^lo^Nur77^-^ cDC1” which are similar to CD207^lo^Nur77^+^ cDC1s, but lack expression of genes including *Nr4a1* (Nur77), *Il1b*, and *Fos*, and “Traf1^+^ cDC1s”, which express elevated *Traf1,* which is critical for NF-kB signaling^45^ (Fig. 5b and Extended Data Fig. 6c). Consistent with flow cytometry data (Extended Data Fig. 4e), cDC1s decline in CD4SP-deficient *MHC-II^-/-^* scRNA-seq datasets (Extended Data Fig. 6f). Notably, the frequency of CD207^hi^Nur77^-^ cDC1s decreases in datasets from *B2m^-/-^* and *Rag2^-/-^* mice, counterbalanced by expansion of CD207^lo^ and Traf1^+^ cDC1 subsets (Fig. 5d,e), indicating CD8SPs alter cDC1 homeostasis beyond what was detectable by flow cytometry (Fig. 3c and Extended Data Fig. 4c).

Three cDC2 clusters matched those detected in WT cDCs (Fig. 1a): EpCAM^+^ cDC2, cycling cDC2, and tDC2 (Fig. 5b,c). Two cDC2 clusters mainly emerged from *Rag2^-/-^*thymi (Fig. 5d,f): “Clec4^+^ cDC2”, with high expression of *Lyz2* and *Clec4a2*, and “Nur77^+^ cDC2”, which transcriptionally overlapped with Nur77^+^ cDC1s (Fig. 5b and Extended Data Fig. 6c). We also identified an intermediate cDC2 (“intDC2”) subset, likely in the process of activation, as it expresses genes characteristic of non-activated (e.g., *Ltb, Sirpa*) and activated (e.g *Ccr7*, *Cd63*) cDC2s (Fig. 5b and Extended Data Fig. 6c). Consistent with CD8SP support of cDC2 homeostasis, cDC2 frequencies decline in *Β2m^-/-^* datasets (Extended Data Fig. 6f), driven by a reduced proportion of tDC2s (Fig. 5f). EpCAM^+^ cDC2s declined substantially in *Rag2^-/-^* thymi, counterbalanced by expansion of Clec4^+^ cDC2s (Fig. 5f), reflecting a larger transcriptional shift in cDC2s than revealed by flow cytometry.

We detected aDC1s, aDC2s and a novel “Il15ra^hi^ aDC” cluster with lower *Ccr7* and *Cd63* expression, which was primarily derived from KO strains (Fig. 5b,d,g). Though flow cytometry revealed greatly diminished aDC numbers in *Rag2^-/-^*thymi (Fig. 3a), scRNA-seq identified a substantial pool of aDCs in *Rag2^-/-^* thymi (Fig. 5d and Extended Data Fig. 6f) suggesting molecular cues independent of DP and SP thymocytes can contribute to cDC activation.

In summary, scRNA-seq data of thymic DCs from *Β2m^-/-^*, *MHC-II^-/-^*, and *Rag2^-/-^*mice corroborate that different thymocyte subsets support homeostasis and activation of distinct DC subsets and reveal additional transcriptional changes not apparent by flow cytometry.

### CD4SP and CD8SP thymocytes promote DC homeostasis and activation via distinct pathways

To determine how SP thymocytes alter transcriptional programs associated with cDC activation, we used p-Creode^46^, an unsupervised algorithm for single-cell trajectory analysis, to reconstruct known differentiation trajectories of WT cDC1s and cDC2s for the purpose of identifying gene programs exhibiting differential dynamics during cDC activation. RNA velocity analysis validated that cDC1 and cDC2 differentiate into their activated counterparts, and CytoTRACE scoring^47^ confirmed reasonable p-Creode mapping with immature cDCs exhibiting high progenitor activity (high scores), and activated DCs exhibiting low differentiation potential (low scores) (Extended Data Fig. 7a,b). While we did not use trajectory reconstruction to define new lineage relationships, the resulting trajectories were consistent with the established intrathymic differentiation from non-activated to activated cDC subsets^8, 9^. To identify the transcriptional states along these trajectories, we assessed the proportion of UMAP-based cDC clusters in each node (Fig. 6a,b).

**Figure 6.**
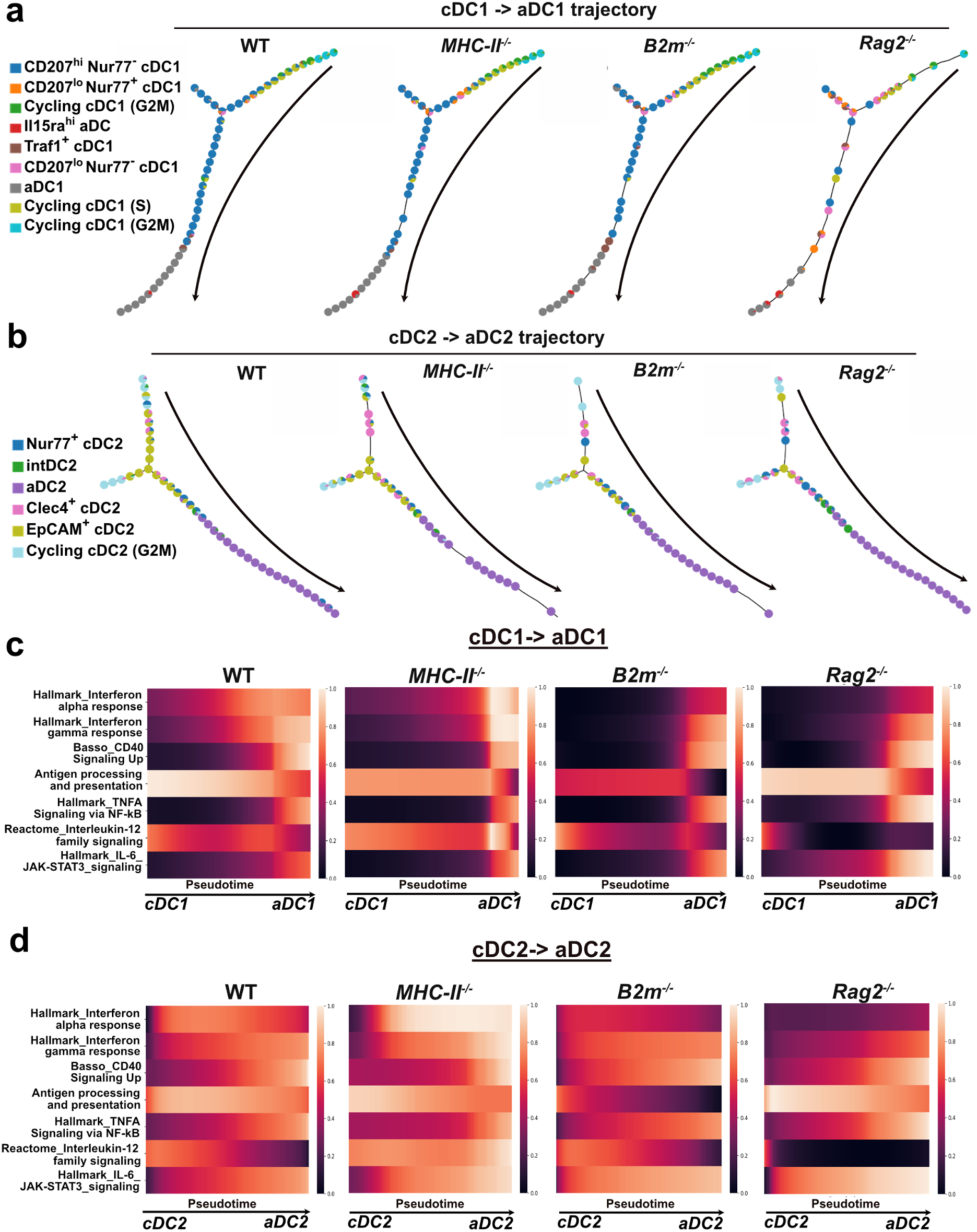
SP thymocytes contribute to DC homeostasis and activation by regulating interferon signaling and NF-kB signaling. (a-b) p-Creode topology depicting transcriptional states (represented as nodes) along the (a) cDC1 and (b) cDC2 differentiation trajectories. The colors in each node represent the proportion of cDC1 and cDC2 transcriptional clusters from Fig. 5a comprising each respective transcriptional state. (c-d) Pseudotime heatmaps displaying signature scores of pathways associated with genes that were differentially expressed along the (c) cDC1– aDC1 and (d) cDC2-aDC2 differentiation trajectories for each WT versus KO comparison. See also Extended Data Fig. 7 and 8.

To determine how thymocyte subsets impact cDC differentiation, scRNA-seq data from KO DCs were projected onto WT p-Creode trajectories. Compared to WT cDC1 and cDC2 trajectories, some differentiation nodes were not populated or were comprised of altered transcriptional subsets along the transition to activated DC states in *MHC-II^-/-^, Β2m^-/-^,* and *Rag2^-/-^* mice, indicating dysregulated cDC differentiation (Fig.6a,b). The unexpected presence of aDC1s and aDC2s in *Rag2^-/-^* scRNA-seq datasets indicates that cDCs undergo at least partial activation without crosstalk from SP thymocytes (Fig. 6a,b). However, *Rag2^-/-^* aDCs express lower levels of genes associated with cDC activation (Extended Data Fig. 7c,d), likely explaining why *Rag2^-/-^*aDCs are not detected by flow cytometry, and underscoring the need for “licensing” from thymocytes to support upregulation of MHC and costimulatory molecules^48^. The replacement of WT-like aDC1 nodes by Il15ra^hi^ aDCs (Fig. 6a) in all KO strains also suggests alternate cDC activation signals in the absence of SP thymocytes.

To elucidate how SP thymocytes support thymic DC activation, we identified differentially expressed genes along the cDC1 and cDC2 activation trajectories between WT and KO genotypes using tradeSeq^49^ and then performed pathway enrichment analysis (Fig. 6c,d). In *MHC-II^-/-^*mice, CD4SP deficiency resulted in upregulation of genes associated with IFNα (Type I) and IFNγ (Type II) signaling along the trajectory of cDC1 and cDC2 activation; conversely, genes associated with IFNα and IFNγ signaling were downregulated along these trajectories in *Β2m^-/-^* and *Rag2*^-/-^ mice (Fig. 6c,d and Extended Data Fig. 8a,b), suggesting CD8SP thymocytes promote IFN signaling in differentiating thymic cDCs. In *MHC-II^-/-^* mice, genes associated with CD40 signaling were downregulated along the cDC1 trajectory, indicating CD4SP thymocytes support cDC1 activation through CD40-dependent non-canonical NF-kB signaling (Fig. 6c).

The absence of SP thymocytes in *Rag2^-/-^* mice results in increased canonical NF-kB and IL-6-JAK/STAT3 signaling, along with a stark decline in IL-12 signaling in aDC1s and aDC2s, further indicating an altered transcriptional state in aDCs that persist without crosstalk from SP thymocytes (Fig. 6c,d and Extended Data Fig. 8c). Altogether, analysis of differentially regulated gene expression along the trajectory of cDC1 and cDC2 activation indicates that IFN signaling, which requires CD8SP cells, and NF-kB signaling, which requires CD4SP cells, cooperate to support cDC homeostasis and activation in the thymus.

### CD8SP thymocytes indirectly regulate pDC survival and tDC2 recruitment to the thymus, and promote Type III IFN production by mTECs

We evaluated mechanisms by which CD8SP cells could regulate pDC and tDC2 homeostasis and IFN signaling during DC activation. Differentiated cDC2s and pDCs migrate into the thymus, where they mediate central tolerance^5, 15, 16, 40^. PSGL-1 contributes to thymic homing of peripheral DCs^40^, and PSGL-1 expression was reduced in thymic pDCs from *B2m^-/-^* mice (Fig. 7a-c). Thus, to test whether *B2m*^-/-^ pDCs and/or cDC2 subsets migrate less efficiently into the thymus, we transferred a mixture of splenic DCs from WT and *B2m^-/-^* mice into non-irradiated congenic WT hosts (Fig. 7d). After 48 hours, the transferred *B2m*^-/-^ pDCs and tDC2s had migrated into host thymi as efficiently as their WT counterparts (Fig. 7d). Thus, despite lower expression of PSGL-1, the decline in pDCs and tDC2s in *B2m^-/-^* thymi is not due to a cell-intrinsic migration defect.

**Figure 7.**
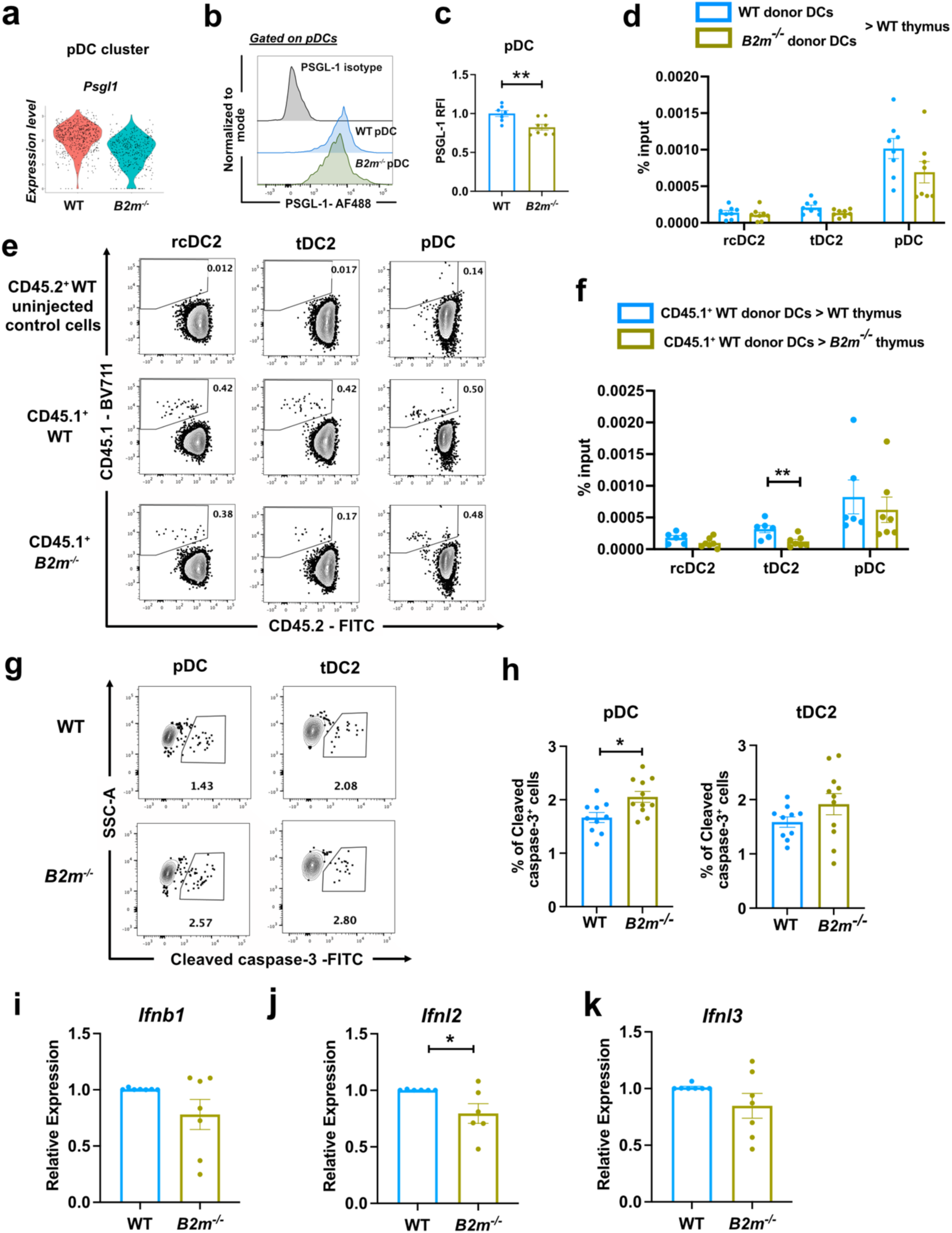
CD8SP thymocytes indirectly regulate intrathymic pDC survival and tDC2 recruitment to the thymus, and promote Type III interferon expression by mTECs. (a) Violin plots showing expression of *Psgl1* transcripts in WT vs *B2m^-/-^*pDCs. (b) Representative flow cytometry histograms showing PSGL-1 expression by thymic pDCs of the indicated genotypes, as well as isotype control levels. (c) Quantification of RFIs of PSGL-1 cell surface expression on thymic pDCs. Data are normalized to average PSGL-1 expression on WT pDCs. Data are compiled from N=3 independent experiments (n=7-8 mice). (d) Quantification of the percentage of donor DCs of the indicated genotypes detected in WT thymi 48 hours after a 1:1 mixture of CD45.2^+^ WT and CD45.2^+^ *B2m*^-/-^ splenic DCs were retro-orbitally injected into non-irradiated CD45.1^+^ congenic WT recipients. Data are from N=3 independent experiments (n=8 mice). (c-d) Bars represent mean ± SEM and symbols represent individual mice. Percentages are calculated relative to the number of DC subsets of each respective genotype transferred. (e) Representative flow cytometry plots and (f) quantification of the percentages of donor-derived thymic DC subsets recovered from recipient mice 48 hours after transfer of CD45.1^+^ WT splenic DCs into either CD45.2^+^ WT or CD45.2^+^ *B2m*^-/-^ recipient mice as calculated in (d). Un-injected CD45.2^+^ WT mice were used as a negative control to set flow cytometry gating. Data are from N=3 independent experiments (n=6-7 mice). Bars represent mean ± SEM and symbols represent individual mice. (g) Representative flow cytometry plots and (h) quantification of the frequency of cleaved caspase-3^+^ pDCs and tDC2s, as indicated, in WT and *B2m*^-/-^ thymi. Data are compiled from N= 5 independent experiments (n= 10-11 mice). Bars represent mean ± SEM and symbols represent individual mice. (i-k) qRT-PCR analysis to quantify transcript expression levels of (i) *Ifnb1*, (j) *Ifnl2*, and (k) *Ifnl3* by WT versus *B2m^-/-^* mTEC^hi^ cells. Bars represent mean ± SEM and symbols represent the average of technical triplicate wells, with cDNA from FACS-purified mTEC^hi^ cells. Data compiled from N=5 independent experiments (n=6-7 pooled mice per experiment). (c, d, f, h, i-k); Statistical analysis was performed using unpaired Student *t*-test *p<0.05, ** p<0.01, ***p<0.001, ****p<0.0001.

We next tested whether the *B2m*-deficient thymic environment fails to efficiently recruit DC subsets from the periphery. 48 hours after injection of WT splenic DCs into non-irradiated mice, a comparable frequency of transferred pDCs migrated into WT versus *B2m*^-/-^ thymi; however, tDC2s were recruited less efficiently into *B2m^-/-^* thymi, highlighting the requirement for MHC-I expression by the thymic environment to maintain tDC2 homeostasis (Fig. 7e,f).

Less than 10% of thymic cDC2s are replaced by circulating DCs in a month, indicating that most cDC2s differentiate within the thymus or are retained therein for prolonged periods^5, 11^. The presence of cycling cDC2s in the thymus (Fig. 5c,f) is also consistent with intrathymic differentiation. Thus, we considered whether altered intrathymic mechanisms contribute to impaired pDC or tDC2 homeostasis in *B2m^-/-^* thymi. The frequencies of proliferating cDC2 subsets and apoptotic tDC2s did not differ in *B2m^-/-^* versus WT thymi (Fig. 5f and Fig. 7g,h), implicating impaired recruitment as the major mechanism underlying fewer tDC2s in *B2m^-/-^* thymi. In contrast, the frequency of pDCs undergoing apoptosis increased in *B2m^-/-^*versus WT thymi (Fig. 7g,h), indicating that CD8SP thymocytes support intrathymic pDC survival.

Data from reciprocal BM chimeras and DC migration assays indicated that CD8SP cells indirectly regulate thymic DC homeostasis and activation by modulating the thymus environment (Fig. 4c-f and Fig. 7d-f). AIRE^+^ mTECs secrete Type I and Type III IFNs, which cooperate to activate HAPCs, including cDC1s^19, 43^. scRNA-seq analyses revealed that Type I IFN response signatures, which overlap with Type III IFN response signatures^50^, decline in thymic DCs in the absence of CD8SP thymocytes (Fig. 6c,d). Hence, to test if CD8SP thymocytes promote IFN production by mTECs, we quantified *Ifnb1*, *Ifnl2*, and *Ifnl3* expression by mTEC^hi^ cells from *B2m^-/-^* versus WT thymi. Notably, expression of *Ifnl2* was reduced in *B2m^-/-^* mTEC^hi^ cells, with similar trends for *Ifnl3* and *Ifnb1* (Fig 7i-k). These data suggest that IFN signaling in thymic DCs, which drives DC activation^19^, is at least in part regulated by CD8SP-mediated support of type III IFN production by mTEC^hi^ cells.

Collectively, these findings suggest CD8SP thymocytes indirectly support pDC survival and tDC2 recruitment to the thymus, and promote Type III IFN production by mTECs to induce IFN signaling in thymic DCs undergoing activation.

### CD4SP thymocytes directly support cDC1 activation via cognate interactions and CD40 signaling, which enforces central tolerance

We next investigated mechanisms by which CD4SP thymocytes promote cDC1 homeostasis and activation. MHC-II expression was required in a cell-intrinsic manner to support cDC1 homeostasis and activation (Fig 4b), and CD40 signaling signatures were reduced along the cDC1 to aDC1 activation trajectory in *MHC-II^-/-^* mice (Fig. 6c). Thus, we tested whether cognate interactions with CD4SP thymocytes and/or CD40 signaling are necessary for cDC1 homeostasis and/or activation.

CD40 is highly expressed by aDC1s and aDC2s in contrast to low expression levels by non-activated cDC subsets (Extended Data Fig. 9a,b). DC-specific CD40 deficiency in *Itgax-cre^+^Cd40^f/-^*mice resulted in a significant decline in thymic aDC1s and a subtle decline in aDC2s, while CD207^+^ cDC1s increased. These findings suggest a block in cDC1 activation at the CD207^+^ cDC1 stage in the absence of CD40 signals (Fig. 8a). Consistent with previous reports^41^, CD4SP thymocytes are the major source of CD40L in the thymus, with CD4SP SM, followed by CD4SP M1 thymocyte subsets^51^ expressing the highest levels of CD40L (Fig. 8b and Extended data Fig. 9c,d). CD40L expression was also detectable on early post-positive selection DP thymocytes (CD3^lo^CD69^+^) and on co-receptor reversing CD3^+^CD69^+^DP cells^27^ (Fig. 8b), indicating that MHC-I- and MHC-II-restricted post-positively selected thymocytes have the potential to activate CD40 signaling in cDCs.

**Figure 8.**
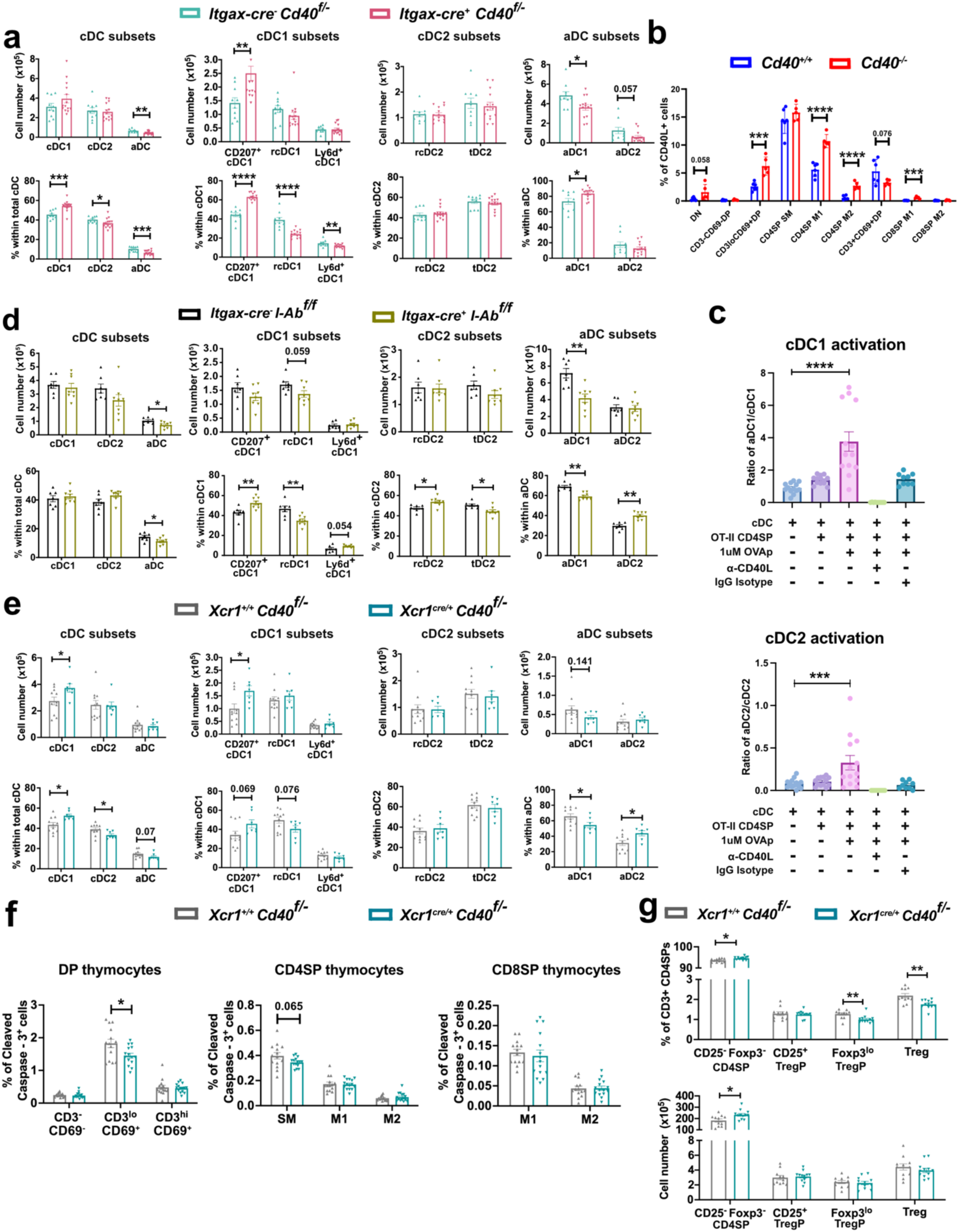
Cognate interactions with CD4SP thymocytes promote CD40 signaling required for cDC1 activation, which is required for central tolerance. (a) Quantification of cell numbers and frequencies of the indicated thymic cDC1, cDC2 and aDC subsets in *Itgax-cre^+^Cd40^f/-^*versus *Itgax-cre^-^Cd40^f/f^* or *Itgax-cre^-^Cd40^f/-^*littermate control mice. The two control genotypes were used interchangeably after observing no discernable differences in the DC compartment between the two genotypes. Data are compiled from N=4 independent experiments (n=10-14 mice) (b) Quantification of the frequencies of each thymocyte subset that express CD40L in *Cd40^+/+^* and *Cd40^-/-^*mice. *Cd40*^-/-^ mice were analyzed because the lack of CD40 receptor engagement prevents CD40L internalization, resulting in maximum detection. (c) Ratio of activated to non-activated cDC1s and cDC2s detected by flow cytometry 17-19 hours after culture of non-activated CCR7^-^CD63^-^MHC-II^lo^ DCs in the presence or absence of OT-II CD4SPs, OVAp_323-339_, and blocking antibodies to CD40L or an isotype control antibody, as indicated. Data are compiled from N=4 independent experiments. Bars represent mean ± SEM and each symbol represents data from one well. (d-e) Quantification of cell numbers and frequencies of the indicated cDC1, cDC2, and aDC subsets in (d) *Itgax-cre^-^I-Ab^f/f^* versus *Itgax-cre^+^I-Ab^f/f^* littermate mice and (e) *Xcr1^+/+^Cd40^f/-^*versus *Xcr1^cre/+^Cd40^f/-^* littermate mice. Data are compiled from N=4 independent experiments (n= 9-11 mice). (f) Quantification of frequencies of cleaved caspase-3^+^ cells within (left to right) DP, CD4SP and CD8SP subsets in *Xcr1^+/+^Cd40^f/-^* versus *Xcr1^cre/+^Cd40^f/-^*littermate mice. Data are compiled from N=5 independent experiments (n=14-15 mice). (g) Quantification of frequencies and numbers of CD25^-^Foxp3^-^ CD4SP conventional thymocytes, CD25^+^Foxp3^-^ TregP cell, CD25^-^Foxp3^lo^ TregP cells, and CD25^+^Foxp3^+^ Tregs in *Xcr1^+/+^Cd40^f/-^*versus *Xcr1^cre/+^Cd40^f/-^* thymi. Data are compiled from N=4 independent experiments (n=11-12 mice). (a-b, d-g) Bars represent mean ± SEM and symbols represent individual mice. Statistical analysis was performed (c) using one-way ANOVA with Dunnett’s multiple comparisons test relative to respective cDC only control and (a, b, d-g) unpaired Student *t*-test; *p<0.05, ** p<0.01, ***p<0.001, ****p<0.0001. See also Extended Data Fig. 9.

To test whether cognate interactions with CD4SP cells induce cDC activation in a CD40-dependent manner, we employed in vitro thymic DC activation assays (Fig. 4g). Once again, only cognate interactions with CD4SPs induced cDC1 and cDC2 activation (Fig. 8c). Addition of anti-CD40L blocking antibodies inhibited cDC1 and cDC2 activation, underscoring the requirement for CD40 signaling driven by cognate interactions with CD4SP cells (Fig. 8c).

Although cognate interactions with CD4SP thymocytes were sufficient to activate both cDC1s and cDC2s, cDC1 activation occurred more efficiently in vitro (Fig. 8c). Moreover, only cDC1 activation was impaired in constitutive *MHC-II^-/-^* mice (Fig. 3d) and in *MHC-II^-/-^* thymic DCs from mixed BM chimera recipients (Fig. 4b). Consistent with these findings, analysis of MHC-II-deficient thymic DCs from *Itgax-cre^+^I-Ab^f/f^*mice confirmed that only cDC1s, but not cDC2s, must express MHC-II to become activated in vivo (Fig. 8d). Together, these findings demonstrate that cognate interactions with CD4SP cells are required for activation of cDC1s, but not cDC2s.

To test if CD40-activated aDC1s are required for central tolerance, we analyzed thymocytes from *Xcr1^cre/+^Cd40^f/-^*mice^52^. As in DC-specific *Cd40*-deficient mice, cDC1-specific CD40 deficiency resulted in an expansion of CD207^+^ cDC1s and a diminished frequency of aDC1s, consistent with a block in cDC1 activation (Fig. 8e). In the absence of CD40-activated aDC1s, the frequency of post-positive selection CD3^lo^CD69^+^ DPs undergoing apoptosis declined (Fig. 8f), and CD4SP SM numbers increased (Extended Data Fig. 9e), suggesting aDC1s play a non-redundant role in early-stage negative selection^27, 53^. Frequencies of Tregs and Foxp3^lo^ Treg progenitors (TregP) also declined in *Xcr1^cre/+^Cd40^f/^*^-^ mice (Fig. 8g), indicating that CD40-activated aDC1s contribute to Treg selection. Furthermore, expression levels of proteins associated with Treg function (CTLA-4, GITR, CD25) were reduced on CD25^+^ TregP from *Xcr1^cre/+^Cd40^f/^*^-^ thymi, implicating CD40-activated aDC1s in generation of fully functional TregP (Extended Data Fig. 9f).

Collectively, our findings demonstrate that CD40 signaling drives thymic cDC activation, but only cDC1s must undergo cognate interactions with CD4SP thymocytes to induce CD40-mediated activation. Our data also highlight a non-redundant role for CD40-signaled aDC1s in negative selection and Treg induction.

## DISCUSSION

Though recent studies have documented roles for several DC subsets in thymocyte selection^2, 4, 54^, the differential cellular and molecular crosstalk signals regulating DC homeostasis and/or activation remain largely enigmatic. Here, we identify transcriptionally distinct thymic DC subsets through scRNA-seq at greater resolution than previously described^11, 26, 55^, elucidate their functional abilities, and characterize microenvironmental signals supporting their homeostasis and activation.

Our studies reveal that while both activated thymic DC subsets express high levels of MHC and co-stimulatory molecules, aDC1s and aDC2s differ in the efficiency with which they present mTEC-derived antigens and induce Treg differentiation. Previous studies showed that aDC1s most efficiently acquired proteins from *Aire*^+^ mTECs^10^, and they presented mTEC-derived antigens on MHC-I and MHC-II more efficiently than non-activated cDC1s^8^. Our data extend these findings by showing that relative to all thymic DC subsets, aDC1s, followed by aDC2s, most efficiently acquire and present mTEC-derived antigens on MHC-I and MHC-II. *Cd36*, a scavenger receptor that facilitates acquisition of cell-surface proteins from apoptotic mTECs^56^, is expressed exclusively by cDC1s (Fig. 1b), possibly accounting for the superior presentation of mTEC-derived antigens by aDC1s. cDC2s must use alternative mechanisms to acquire mTEC-derived antigens, possibly via trogocytosis, exosome transfer, or gap junctions^4^. In agreement with reports from our lab and others^7, 34, 57^, cDC2s/aDC2s induced Treg differentiation more efficiently than cDC1s/aDC1s or pDCs. However, aDC1s have been reported to select some TRA-specific Tregs^56, 58^. Whether aDC1s and aDC2s induce tolerance to TRAs versus other self-antigens, respectively, remains unresolved, as do mechanisms underlying the superior ability of DC2/aDC2s to induce Treg.

Few microenvironmental signals that support DC homeostasis and activation within the thymus have been elucidated. The impairment in DC activation in genetic models in which thymocytes are blocked pre-positive selection is consistent with prior studies of *Tcra*^-/-^ thymic DCs^9, 18^. Moreover, cDC1 numbers declined in *Zap70^-/-^* mice^8^, consistent with our data showing that SP thymocytes are required to maintain cDC1 homeostasis. Cognate interactions with CD4SP thymocytes and CD40 signaling were reported to drive thymic cDC activation, while a role for CD8SPs was not identified^9, 18^. In contrast, our data reveal that CD4SP and CD8SP thymocytes play distinct, essential roles in regulating homeostasis and activation of cDC1s versus cDC2s and pDCs, respectively. Such divergent conclusions may reflect the refined gating schemes we used to identify aDCs, informed by scRNAseq analyses. Since thymocytes drive TEC maturation^41, 42, 59^, and TECs produce cytokines that promote DC activation^19, 43^, we tested whether thymocytes directly interacted with DCs or indirectly modified the thymus environment to regulate DC homeostasis and activation. Our findings indicate that while CD4SP thymocytes undergo direct cognate interactions required for DC homeostasis and cDC1 activation, CD8SPs indirectly promote pDC and tDC2 homeostasis by modulating the thymic environment.

Single-cell transcriptional profiling of thymic cDCs from *B2m^-/-^, MHC-II^-/-^* and *Rag2^-/-^* mice suggested that CD8SPs induce IFN signaling to regulate cDC1 and cDC2 activation, while CD4SPs promote cDC1 differentiation and activation through CD40-dependent non-canonical NF-kB signaling. In the context of infections, IFN signaling promotes peripheral DC activation and enhances expression of costimulatory molecules for effective naïve T cell stimulation^60, 61^. In the sterile thymus, mTECs constitutively express Type I and Type III IFNs^19, 43, 62^. While previous studies highlight a critical role for *Aire* in IFN expression by mTECs^19^, we find that CD8SP cells are required for efficient expression of Type III IFNs by mTEC^hi^ cells. In *Ifnar*^-/-^*Ifnlr^-/-^* mice, IFN signaling declines in thymic DCs, cDC1 activation is impaired, and Treg TCR diversity is reduced^19^. Thus, reduced IFN expression by mTEC^hi^ cells in *B2m*^-/-^ thymi likely contributes to the diminished Type I IFN response signature observed in cDC1s and cDC2s, likely impairing cDC activation in the absence of CD8SP thymocytes. Moreover, as Type III IFNs enhance pDC survival in tumor microenvironments^63^, the decline in IFN production may also contribute to reduced pDC survival in *B2m*^-/-^ thymi. Several chemokine receptors have been implicated in recruitment of DCs to the thymus. For instance, some cDC2s require CCR2 and CX3CR1 for thymic localization, while CCR2 and CCR9 mediate pDC recruitment to the thymus^15–17^. We found that tDC2s migrated less efficiently into *B2m^-/-^* thymi, suggesting CD8SPs may also regulate expression of chemokines by thymic stromal cells.

We tested the hypothesis, suggested by scRNA-seq analyses, that CD40 is required for cDC1 homeostasis and activation. Using a novel in vitro thymic DC activation assay, we found that CD4SP-derived cognate interactions were sufficient to activate thymic cDC1s and cDC2 in a CD40-dependent manner. These findings are consistent with prior reports that cognate interactions with CD4SPs can activate BM-derived DCs in a CD40-dependent manner^9^ and that thymic DCs can be activated by injection of cognate peptides into TCR transgenic mice^18^. However, while CD40 expression was required for efficient DC2 activation, antigen presentation on MHC-II was necessary only for CD40-dependent activation of cDC1s in vivo. Thus, although we and others find that CD4SP thymocytes are the major source of CD40L in the thymus^41^, other cellular sources of CD40L, such as iNKTs which have been shown to induce IL-12 production in cDC1s through CD40-CD40L interactions^64^, DN thymocytes, or DP thymocytes selected on MHC-I (Fig. 8b), could redundantly support thymic cDC2 activation.

Though several studies have demonstrated the role of aDC1s in mTEC-derived antigen acquisition, their role in central tolerance remains contentious^56–58, 65^. Our findings highlight a non-redundant role for CD40-activated aDC1s in both early stages of negative selection and efficient Treg induction, consistent with a recent study showing that a reduction in aDC1s correlates with a decline in TregP and Treg cells^19^. Moreover, in line with our data showing that aDC1s promote negative selection, cDC1-deficiency led to selection of T cells that caused multi-organ autoimmunity when transplanted into immunodeficient mice^56^. Together, our findings and prior studies reveal that aDC1s are required for mTEC-derived TRA acquisition and efficient central tolerance.

Future studies will test candidate mechanisms by which CD8SP cells indirectly regulate the thymus environment to support IFN signaling and pDC and tDC2 homeostasis, and will elucidate the relative contributions of cDC1 versus cDC2 subsets to central tolerance against distinct classes of self-antigens. Understanding mechanisms of thymocyte-stromal-DC crosstalk will provide a framework for altering the thymic DC activation state to efficiently promote selection of a self-tolerant T cell pool.

## METHODS

### Mice

C57BL/6J (Wild-type; WT; B6), BALB/cJ, B6.SJL-Ptprc^a^PepC^b^ (CD45.1), C57BL/6J x B6.SJL-Ptprc^a^PepC^b^ (F1), C57BL/6-Tg(TcraTcrb)1100Mjb/J (OT-I), B6.Cg-Tg(TcraTcrb)425Cbn/J (OT-II), C57BL/6-Tg(Ins2-TFRC/OVA) 296Wehi/WehiJ (RIP-mOVA) ^30^, RIP-OVA^hi^ ^31^ (W. R. Heath, University of Melbourne, Melbourne, Australia), B6(Cg)-Rag2^tm1.1Cgn^/J (*Rag2^-/-^*), B6.129S2-H2^dlAb1-Ea^/J (*MHC-II^-/-^*), B6.129P2-B2m^tm1Unc^/DcrJ (*B2m^-/-^*), B6.129S6-Del(3Cd1d2-Cd1d1)1Sbp/J (*Cd1d^-/-^*), B6.129S2-Tcra^tm1Mom^/J (*Tcra^-/-^*), B6.Cg-Tg(Itgax-cre)1-1Reiz/J (*Itgax-cre^+^*), B6-129X1-*H2-Ab1^tm1Koni^*/J (*I*-*Ab^f/f^*), Foxp3^tm9(EGFP/cre/ERT2)Ayr^/J, *Cd40^f/f^* ^52^, *Xcr1^cre^*^/+^*Cd40^f^*^/-52^, and *Cd40*^-/- 66^ (K. Murphy, Washington University School of Medicine, St. Louis, Missouri, USA) strains were bred in-house. Experiments were performed using 6–8-week-old mice of both sexes. All strains were purchased from The Jackson Laboratory unless otherwise specified. Mice were maintained under specific pathogen-free conditions at the Animal Resources Center, the University of Texas at Austin. All experimental procedures were performed in accordance with the Institutional Animal Care and Use Committee.

### Flow cytometry and FACS

For isolating thymic cDCs and TECs, thymi of the appropriate genotypes were isolated, cut into small pieces, and enzymatically digested as previously described^67^. Briefly, each thymus was digested in 2mL of enzymatic mixture containing Liberase^TM^ (Roche) and DNAse I (Roche) in PBS at 37°C for 12 minutes, gently swirling the tubes mid-interval (6 minutes). After digestion, supernatant containing cells released from the tissue was transferred into 30mL of FACS-wash buffer (FWB)+ EDTA (PBS+ 2% bovine calf serum (BCS) + 5mM EDTA) and stored on ice. The enzymatic digestion step was repeated twice on residual tissue fragments to ensure complete dissociation of the tissue. Digested cells were centrifuged in 2mL of FWB+EDTA at 300g for 5 minutes at 4°C, the supernatant was discarded (henceforth referred to as “washing the cells”), and the cell pellet was resuspended in FWB+ EDTA. For thymic DC flow cytometry panels that included anti-CCR7, 6 x 10^6^ cells were stained with the indicated cocktail of fluorophore conjugated-antibodies in 200µL FWB for 40 minutes at 37°C before washing the cells once and resuspending in FWB+EDTA with 0.1µg/mL propidium iodide (Enzo).

For thymocyte staining panels that included anti-Foxp3, thymocyte single cell suspensions were generated by pressing thymi through 40µm cell strainers, which were rinsed with FWB. 6 x 10^6^ cells were stained with the appropriate cocktail of cell-surface conjugated antibodies in 200µL FWB with the fixable viability dye Zombie Red (BioLegend; 1:1000) for 30 minutes on ice. After washing the cells once in FWB and resuspending the cells in 200µL of FWB (PBS + 2% BCS), cells were fixed and permeabilized with the Foxp3 Transcription factor staining Buffer kit according to manufacturer’s recommendations (Tonbo Biosciences), followed by intracellular staining with anti-Foxp3-PE (ThermoFisher) for 30 minutes on ice. To FACS-sort CD4SP thymocytes, after antibody staining, cells were washed with FWB and resuspended in FWB plus propidium iodide (Enzo), as above. For functional assays, cells were FACS-sorted into PBS+ 10% BCS.

To quantify apoptosis, using cleaved caspase-3 staining, single cell suspensions were first generated by passing the thymi through 40µm cell strainers, which were rinsed with FWB. 6 x 10^6^ cells were stained with the appropriate cocktail of cell-surface conjugated antibodies in 200µL FWB and the fixable viability dye Zombie Red (BioLegend; 1:1000) for 30 minutes on ice. After washing once in FWB and resuspending the cells in 200µL of FWB (PBS + 2% BCS), cells were fixed and permeabilized with the BDCytofix/Cytoperm kit according to manufacturer’s instructions prior to incubation with anti-cleaved caspase-3 for 30 minutes on ice as previously described^53^. Cells were then washed and resuspended in FWB before flow cytometry analysis.

To quantify CD40L expression (Fig. 8b and Extended Data Fig. 9c,d), thymocytes from WT mice were isolated and stained for cell surface expression of CD3, CD4, CD8, CD69, MHC-I, CD73, and non-αβT-cell lineage antibodies (CD19, B220, NK1.1, Gr-1, TCRγδ, Ter119, CD11c, CD11b). The cells were then fixed with BD Cytofix/Cytoperm as per the manufacturer’s instructions. WT thymocytes were then stained for intracellular CD40L or for an isotype control, while thymocytes isolated from *Cd40^-/-^* mice were stained only for surface expression of CD40L without fixation.

For flow cytometry analysis or FACS-isolation of the indicated cell types, cocktails of the following fluorescent-conjugated antibodies were used-CD11b (M1/70), CD11c- (N418), SIRPα (P84), CX3CR1 (SA011F11), XCR1 (ZET), F4/80 (BM8), Ly6d (49-H4), PDCA-1 (927), CD14 (M14-23), CD207 (4C7), CCR7 (4B12), CD63 (NVG-2),I-A/I-E (M5/114.15.2), CD19 (1D3), CD90.2 (53-2.1), CD45 (30-F11), Y-Ae (eBioY-Ae), Vα2 (B20.1), Vβ5 (MR9-4), TCRβ (H57-597), CD3 (145–2C11), CD45.1 (A20), CD80 (16-10A1), CD86 (A17199A), CD40 (FGK45), H2-Kb (AF6-88.5), CD4 (RM4-5), CD8 (53-6.7), CD25 (PC61), FOXP3 (FJK-16s), CD73 (TY/11.8), EpCAM (G8.8), CD69 (H1.2F3), CD40L (MR1), NK1.1 (PK136), Gr-1 (RB6-8C5), CD19 (6D5), B220 (RA3-6B2), TCRγδ (eBioGL3), Ter119 (TER-119), CD45.2 (104), CD80 (16-10A1), Ly51 (6C3), Helios (22F6), PD-1 (29F.1A12), GITR (YGITR 765), CD5 (53-7.3), CTLA-4 (UC10-4B9), Icos (7E.17G9), CD44 (IM7), Cleaved Caspase-3 (D3E9). All antibodies were obtained from BioLegend, eBioscience, or BD Biosciences. The lectin UEA-1 (Vector Laboratories) was used to identify mTECs. All antibodies and lectins were used at a 1:400 dilution to stain the respective cell types (6 x 10^6^ cells in 200 µL volume) except as noted: CCR7 (1:200), I-A/I-E (1:5000), FOXP3 (1:200), CD40L (1:200), NK1.1 (1:800), CD19 (1:800), B220 (1:800), TCRγδ (1:800), Ter119 (1:800), CD11c (1:800), CD11b (1:800), and EpCAM (1:800).

Flow cytometry analysis was carried out on a BD LSRFortessa^TM^ or a BD FACSAria^TM^ Fusion. All FACS-sorting was carried out on a BD FACSAria^TM^ Fusion. Flow cytometry data were analyzed using FlowJo software (V10.8.0, BD Biosciences).

### FACS-isolation of cDC subsets for functional assays

Thymi from 5 mice of the appropriate genotypes were extracted for the corresponding assays and enzymatically digested as described above to isolate thymic DCs. Prior to FACS-sorting DC subsets, a DC-enrichment step was carried out: after washing the cells in FWB+EDTA following enzymatic digestion, they were washed and stained with CD11c-Biotin (N418, Biolegend) for 20 minutes at 4°C. After washing once in FWB+EDTA, the cells were incubated with 10µL of Streptavidin Microbeads (Miltenyi Biotec) per 10^7^ cells for 15 minutes on ice and then washed once with FWB+EDTA. CD11c-stained cells were positively enriched using MACS LS columns in the magnetic field of a MidiMACS separator according to the manufacturer’s protocol. CD11c-enriched cells were then stained with antibodies against CCR7, CD11b, CD11c, CD14, CD63, CD207, CX3CR1, F4/80, Ly6d, MHC-II, PDCA-1, SIRPα, and XCR1 for FACS-sorting the 8 DC subsets –pDCs (SSC-A^lo^PDCA1^+^CD11c^+^), CD207^+^ cDC1s, Ly6d^+^ cDC1s, rcDC1s (Ly6d^-^CD207^-^), tDC2s (CD14^+^ and/or CX3CR1^+^), rcDC2s (CD14^-^CX3CR1^-^), aDC1s (CCR7^+^CD63^+^XCR1^+^), and aDC2s (CCR7^+^CD63^+^SIRPα^+^) - using the gating strategy shown in Extended Data Fig. 2a.The 8 DC subsets were sorted into PBS+ 10% BCS using a 100µm nozzle on the BD FACSAria^TM^ Fusion.

### In vitro OT-I DC stimulation assays

Thymic DC subsets were FACS-sorted as described above from RIP-mOVA and RIP-OVA^hi^ mice and resuspended at a cell concentration of 10,000 cells/100µL in complete RPMI (RPMI 1640 (Gibco), 10% fetal bovine serum (FBS; Gemini Bio-Products), 1 mM Sodium Pyruvate (Gibco), 1x GlutaMAX (2 mM L-alanyl-L-glutamine dipeptide; Gibco), 1x MEM Non-essential Amino Acid Solution (Sigma-Aldrich), 1x Penicillin-Streptomycin-Glutamine (100 units of penicillin, 100 µg of streptomycin, and 292 µg/ml of L-glutamine; Gibco), and 55 µM β-mercaptoethanol (Gibco)). CD8^+^ T cells were enriched from the spleen of OT-I mice using the Mojosort^TM^ Mouse CD8^+^ T cell isolation kit (Biolegend) and stained with 5µM CellTrace Violet (Thermo Fisher) according to manufacturers’ instructions. Enriched CellTrace Violet-labeled OT-I CD8^+^ T cells were resuspended at 20,000 cells/100µL in complete RPMI + 10ng/mL of recombinant mouse IL-2 (Biolegend). As positive control, WT splenocytes were pulsed with 50nM exogenous OVA_257-264_ (SIINFEKL) peptide (New England Peptide) for 30 minutes at 37°C and then washed once with complete RPMI following incubation. cDC subsets from the RIP-mOVA or RIP-OVA^hi^ mice or OVA_257-264_ -pulsed WT splenocytes were cocultured with CellTrace Violet-labeled OT-I CD8^+^ T cells at a 1:2 ratio for 3.5 days at 37°C for before CD8^+^ T cell proliferation was assessed by flow cytometry. The number of cell divisions was determined using the proliferation modeling tool in FlowJo^TM^ software. Unstimulated CD8^+^ T cells were used to identify “Generation 0” peak, and a peak ratio close to 0.5 percent was implemented, indicating that every cell division results in half as much dye in the cells from the next generation. Each proliferation peak was gated by the tool to determine the percentage and number of cells associated with distinct rounds of cell division.

### In vitro Treg generation assays

Treg generation assays were set up as previously described^34^. 7 cDC subsets and 1 pDC subset were FACS-sorted as discussed above from 4-5 pooled B6 thymi and each subset was reconstituted at a concentration of 10,000 cells/100µL in complete RPMI. For Treg generation assays with cDCs isolated from T-cell subset-deficient mice, thymi from WT, *MHC-II^-/-^*, *B2m^-/-^*, and *Rag2^-/-^*were enzymatically digested and enriched for DCs as described above. DCs were then FACS-purified into cDC1s, cDC2s, aDC1s, and aDC2s as shown in Extended Data Fig. 2a.

CD73^-^CD4^+^CD25^-^ Foxp3^-^ newly generated CD4SP thymocytes were sorted from *Foxp3^eGFP-Cre-ERT2^* reporter mice and reconstituted at a concentration of 20,000 cells/100µL in complete RPMI + 100ng/mL recombinant mouse IL-7 (Biolegend). CD4SP thymocytes and DC subsets were co-cultured at a 2:1 ratio in round-bottom 96-well plates for 4 days and then analyzed by flow cytometry on day 5 to evaluate the percentage of Tregs generated in vitro.

### FACS purification of thymic HAPC and cDC subsets for scRNA-sequencing

To sort HAPCs for scRNA-sequencing, thymi were harvested and enzymatically digested as described above. After staining with a cocktail of antibodies, as shown in Extended Data Fig. 1a, cells were resuspended in FWB+EDTA plus propidium iodide (Enzo) for FACS-sorting. We FACS-isolated CD45^+^MHC-II^+^Thy1.2^-^ thymic HAPCs from 1 month-old C57BL/6J, *B2m^-/-^* and *Rag2^-/-^* thymi (Extended Data Fig. 1a). From *MHC-II*^-/-^ thymi, we gated on all CD45^+^ cells and then FACS-sorted HAPCs after gating out TECs and thymocytes as in Extended Data Fig. 1a. Sorted cells were washed into FWB without EDTA, counted and resuspended at a concentration of 700-1200 cells/µL prior to submission to The University of Texas at Austin’s Genomic Sequencing and Analysis Facility (GSAF) core.

For hashtagging FACS-isolated cDC-subsets, CD11c^+^ cells were enriched with MACS Streptavidin microbeads, as described above, and stained with the cell surface antibody sorting panel used to delineate cDC subsets, as indicated in Extended Data Fig. 2a. After the 7 cDC subsets were FACS-sorted, each was incubated with a distinct TotalseqB Hashtag antibody (Biolegend) at 0.05µg/10^6^ cells concentration for 30 minutes at 4°C. After staining with the Hashtag antibodies, the cells were washed four times in FWB before pooling 10,000 cells of each sorted DC population, with the exception of Ly6d^+^ cDC1s for which there were fewer remaining cells than the other 6 populations at the end of the protocol. Cells were spun down in FWB without EDTA and resuspended at a concentration of 700-1200 cells/µL prior to sample submission for scRNA-sequencing to the GSAF core.

### scRNA-seq library preparation

Single cell suspensions were processed for scRNA-seq at UT Austin GSAF. Cell suspensions were loaded on a Chromium Controller (10X Genomics) and processed for cDNA library generation following the manufacturer’s instructions for the Chromium NextGEM Single Cell 3’ Reagent Kit v3.1 (10X Genomics). For hashtagged FACS-sorted DC samples, cell surface protein libraries were constructed using the Chromium NextGEM Single Cell 3’ Reagent Kit v3.1 with Feature Barcoding technology (10X Genomics). The resulting libraries were examined for size and quality using the Bioanalyzer High Sensitivity DNA Kit (Agilent) and cDNA concentrations were measured using the KAPA SYBR Fast qPCR kit (Roche). Samples were sequenced on an Illumina NovaSeq 6,000 instrument (paired end, read 1: 28 cycles, read 2: 90 cycles) with a targeted depth of 50,000 reads/cell.

### scRNA-seq preprocessing, visualization, and clustering

In total, 8 thymus HAPC datasets were generated from 4 genotypes: Wild-type (WT) (n=2), *B2m^-/-^* (n=2), *MHC-II^-/-^* (n=2), and *Rag2^-/-^* (n=2). Multiple cell clustering analyses were conducted using combinations of the samples and subsets of cells-of-interest (see below). scRNA-seq FASTQ reads were aligned to the C57BL/6J mm10 reference genome and quantified with Cell Ranger count version 6.0.1.^68^. Cell clustering and visualization analyses were performed using Seurat version 4.1.1^69^ in R version 4.2.2 (R Core Team, 2022).

We first analyzed WT HAPC scRNA-seq data to characterize the cellular heterogeneity of antigen presenting cells in the thymus of physiologically normal C57Bl/6J mice. Briefly, biological replicate samples were merged, and batch-corrected with ‘FindIntegrationAnchors’ and ‘IntegrateData’ Seurat functions. A minimum feature count/cell of 500, a maximum feature count/cell of 7000, and maximum mitochondrial content of 20% were specified as cutoffs. Next, we integrated all samples (n=8) to characterize cellular heterogeneity across genotypes. For WT and KO mice, samples were merged but not batch-corrected because variables-of-interest were confounded with sequencing batch. A minimum feature count/cell of 1000, a maximum feature count/cell of 7000, and maximum mitochondrial content of 20% were used. For both WT-only and all sample integration analyses, cDCs were subset and re-analyzed separately from other cell types to characterize additional heterogeneity within the cDC compartment.

Expression counts were normalized using the ‘NormalizeData’ function and the top 2000 most variable genes were determined with ‘FindVariableFeatures.’ We then performed principal components analysis and used the top 30 principal components as input for k-nearest neighbor mapping and clustering using the ‘FindNeighbors’ and ‘FindClusters’ functions respectively. A resolution of 0.8 was used for clustering datasets initially, but a resolution of 1.5 was used to generate high resolution clusters for the merged samples. Finally, cells were projected into a UMAP dimension reduction with the ‘RunUMAP’ function.

For the WT hashtagged cDC dataset, FASTQ reads were aligned to the mm10 reference genome and quantified with Cell Ranger using the workflow for TotalSeqB samples^68^. Cell clustering and visualization analyses were performed using Seurat version 4.1.1^69^ in R version 4.2.2 (R Core Team, 2022). Hashtagged oligo counts were processed with the “HTODemux” function in Seurat to assign cells to their respective subsets. Cells identified as doublets or negative for hashtag sequence counts were removed prior to gene expression analysis. Gene expression counts were then processed as described above with a minimum feature counts/cell of 1500, a maximum counts/cell of 7000, and a maximum mitochondrial content of 10%. Clusters identified from the hashtagged cDC dataset were annotated based on gene expression similarity to the original cDC dataset (Extended Data Fig. 1d).

Gene markers for each cDC cluster were calculated with the “FindAllMarkers’ function, and gene expression analyses were performed with the ‘FindMarkers’ function, using a minimum log-fold change of 0.25 and a minimum of 10% of cells within the cluster expressing the gene. Cell phase was calculated from expression data using the ‘CellCycleScoring’ function with default cell cycle genes provided in Seurat. Cell classes were then manually annotated for the calculated clusters based on curated lists of genes from Immgen database^70^, Gene Expression Commons^71^, and other relevant publications^11, 21, 55^. Cellxgene was used for scRNA-seq data visualization and exploration (https://github.com/chanzuckerberg/cellxgene).

### scRNA-seq downstream analysis

For p-Creode analyses, Velocyto and scVelo (dynamic mode) were performed initially using the 10x Genomics BAM file from conventional dendritic cell (cDC) subsets, as documented previously^72–74^. Directionality converging to two activated cDC states enabled separation of the dataset into two distinct trajectories. p-Creode analysis was then carried out on data subsets as previously described ^46^. Briefly, principal component analysis was conducted for dimension reduction while maintaining linearity in the data, and end states were identified in an unsupervised manner by closeness centrality value. Transition topologies were constructed using hierarchical placement, which iteratively connects end states and path nodes until the graph component is completed. Consensus topologies were then identified through resampled runs. RNA velocity vectors were visualized on top of the same UMAP embedding to delineate directionality of the transition through the topology. Ly6d^+^ cDCs were excluded from the trajectory due to their high transcriptional similarity to pDCs, suggesting a distinct origin^75^. tDC2s were excluded from the cDC2 trajectory because previous studies indicate they migrate to the thymus from the periphery and do not differentiate intrathymically^12, 13^.

To delineate genes whose dynamics are altered between the conditions (WT versus KO genotypes), tradeSeq^49^ was applied to conduct conditionTest, which assesses differential expression patterns of the average gene expression with pairwise comparison between WT and KO cDC1 and cDC2 trajectories. Gene set enrichment analysis^76, 77^ was performed on differentially expressed genes from tradeSeq to identify significantly enriched biological programs and pathways. A gene signature for each pathway was developed using the union lists of genes for pairwise comparisons (WT vs *B2m^-/-^*, WT vs *MHC-II^-/-^,* and WT vs *Rag2^-/-^*) and signature score trends were computed by fitting generalized additive models (GAMs) over pseudotime. These trends were plotted as heatmaps with gene expression values normalized globally for all conditions.

For scRNA-seq correlation analyses with a published dataset, the top 200 gene markers for each cluster in each dataset were determined and combined into a gene list of common markers. Next, we calculated the average expression of all markers per cluster in each dataset and then calculated correlations between clusters using the Spearman rank correlation. Correlations were then visualized as a heatmap using the ‘pheatmap’ package. For the union of genes identified by tradeSeq analysis per WT versus KO comparison for cDC1 and cDC2 trajectories, we performed functional enrichment analysis on the genes using the R package ‘WebGestaltR’ ^78^, and generated heatmaps showing gene expression for pathways of interest for UMAP-based cDC1, aDC1, cDC2, and aDC2 clusters.

### Generation and analysis of Y-Ae BM chimera mice

BM cells were isolated from the long bones of 6-week-old CD45.1 congenic mice. Bones were crushed and single cell suspensions were prepared in FWB, followed by red blood cells (RBC) lysis (RBC lysis buffer, Biolegend). Cells were stained with the following lineage (Lin)-specific antibodies (all antibodies from BioXcell) - anti-CD11b (M1/70), anti-Gr-1 (RB6-8C5), anti-Ter-119 (TER-119), anti-B220 (RA3.3A1/6.1), anti-CD19 (1D3), anti-CD3 (17A2), and anti-CD8 (53.6.72). Lin^+^ cells were depleted using sheep anti-rat IgG immunomagnetic beads (Dynabeads, Invitrogen). 8-10 x 10^6^ Lin-depleted BM cells from CD45.1 congenic mice were retro-orbitally injected into 6-8-week-old lethally irradiated (1200 rads, delivered in 2 split doses separated by three hours) BALB/c and control B6 recipient mice. Mice were maintained on antibiotic feed (Teklad medicated diet, Inotiv, TD 06596) one day prior and 3 weeks after transplantation. Thymic DCs were analyzed by flow cytometry 8 weeks after reconstitution, including analysis of Y-Ae antibody binding to each DC subset.

### Immunofluorescence of thymic sections

Thymi from C57BL/6J, *MHC-II^-/-^*, *B2m^-/-^*and *Rag2^-/-^* mice were snap-frozen in Tissue-Tek OCT (Sakura) and 7µm cryosections were generated with CryoStar™ NX50 cryostat (Thermo Fisher). Tissue sections were fixed in acetone for 20 minutes at -20°C. After fixation, tissue sections were rinsed with PBST thrice for 3 minutes each and then blocked with TNB (PerkinElmer) for 15 minutes at room temperature (RT). To determine CD207^+^ cDC1 localization, slides were stained overnight (O/N) at 4°C with anti-CD207 (929F3.01, DENDRITICS), anti-CD11c-biotin (N418, Biolegend) and anti-XCR1-AF647 (ZET, BioLegend). Thymic slices were washed thrice at RT with PBST. Streptavidin-AF594 (Thermo Fisher) and Donkey anti-rat 488 (Jackson ImmunoResearch) were added for 1 hour at RT. To detect aDC1 or CD14^+^ cDC2 subsets in the thymus, tissue sections were stained O/N 4°C with either anti-CD11c-biotin (N418, Biolegend), anti-CD63 (NVG-2, Biolegend) and anti-XCR1-AF647 (ZET, BioLegend) for aDC1s or anti-CD11c-biotin (N418, Biolegend), anti-SIRPα (P84, Biolegend) and anti-CD14 (ab231852, Abcam) antibodies for CD14^+^ cDC2s. For aDC1 stains, after washing, sections were then stained with Streptavidin-AF488 (Life Technologies) and Donkey anti-rat 594 (Jackson ImmunoResearch) for 1h at RT. For CD14^+^ cDC2, sections were then stained with Streptavidin-AF488 (Life Technologies), Donkey anti-rat 594 (Jackson ImmunoResearch) and Donkey anti-rabbit 647 (Jackson ImmunoResearch) for 1 hour at RT. For staining aDC2 or pDC subsets, tissue sections were stained O/N 4°C with either anti-CD11c (N418, Biolegend) and anti-CD63 (NVG-2, Biolegend) for aDC2s or anti-CD11c (N418, Biolegend) and anti-SiglecH (552, Biolegend) for pDCs. Next, for aDC2 and pDC stains, sections were incubated with donkey anti-rat 594 (Jackson ImmunoResearch) for 1 hour at RT. Sections were then blocked with 2% rat serum at RT for 1 hour. Following blocking, anti-SIRPα-biotin (P84, Biolegend) was added, and slides were incubated for 1 hour at RT for aDC2s. Slides were incubated with anti-B220-biotin (RA3-6B2, Biolegend) for 1 hour at RT for the pDC stain. Next, slides for both aDC2 and pDC stains were washed and incubated with goat anti-armenian hamster 647 (Biolegend) and Streptavidin -AF488 (Life Technologies) for 1 hour at RT. To stain for macrophages, slides were stained with Mertk-APC (D5MMER, Invitrogen), CD63-PE-DazzleRed594 (NVG-2, Biolegend) and SIRPα-Biotin (P84, Biolegend) overnight at 4°C. The next day, slides were washed twice with PBS and once with PBST for 3 minutes each. Lastly, slides were incubated with Streptavidin-AF488 (Life Technologies) for 1 hour at RT.

All slides were washed thrice with PBST before staining with DAPI and washing again. Slides were then mounted in ProLong^TM^ Gold antifade reagent (Thermo Fisher) and imaged with a 20x and 40x objective on a DMi8 microscope (Leica).

### Histocytometry

Histocytometry was based on analytical approaches previously described^79^. The entire thymus lobe was segmented using a Gaussian mixture model based on DAPI expression. Within this resulting region, another Gaussian mixture model using CD11c or CD63, where appropriate, was used to segment the medulla. Regions outside of the medulla were defined as the cortex. Manual adjustments were made in FIJI to remove imaging artifacts and correct misclassified areas. The CMJ region was then defined as the surrounding 50 µm on either side of the cortical-medullary boundary. After subtracting overlapping CMJ from both the medulla and cortex, the thymic regions were encoded one through three to create a unified mask. This mask was concatenated with the original images prior to cell segmentation, allowing the quantified single-cell data to include regional location.

To identify DC subsets and macrophages, a custom cell segmentation model using DAPI and CD11c or DAPI and CD63 fluorochrome expression was pre-trained on Cellpose^80^. Model fine-tuning was done using the Cellpose 2.0 GUI ^81^ and manual training input. Each DC subset was identified by a distinct combination of cell-surface markers (as in Fig. 1e and Extended Data Fig. 2g): CD207^+^ cDC1 (CD11c^+^XCR1^+^CD207^+^), CD14^+^ cDC2 (CD11c^+^SIRPα^+^CD14^+^), aDC1 (CD11c^+^XCR1^+^CD63^+^), aDC2 (CD11c^+^SIRPα^+^CD63^+^), and pDC (CD11c^+^SiglecH^+^B220^+^). For each cell, the median fluorochrome intensities for all channels and median encoded values for the thymic region mask were quantified using MCquant from the MCMICRO pipeline^82^. To ensure quality, cells were required to meet a minimum area threshold of 150 pixels or 71.74 µm^2^. DC subsets were determined by manual gating via FlowJo. For each cDC subset, cellular density within each thymic region was calculated as the ratio of the number of cells in a given region to the area of that region in mm^2^. To correct for macrophages included within identified aDC2s (CD11c^+^SIRPα^+^CD63^+^), the proportion of CD63^+^SIRPα^+^MerTK^-^ cells out of total CD63^+^SIRPα^+^ cells was calculated within each thymic region for three replicates. The median proportion of CD63^+^SIRPα^+^MerTK^-^ cells was multiplied by the initial aDC2 cell counts per thymic region to estimate the true aDC2 population without macrophages. Figures were generated with R 4.3.1 and Python.

### In vitro thymic DC activation assay

Thymi from four WT mice (Fig. 4g and Fig. 8c) and four *Rag2^-/-^* mice (Fig. 4g) were digested to generate single-cell suspensions and were enriched for DCs as described above. Enriched cells were immunostained with antibodies shown in Extended data Fig. 2a at 37°C, and non-activated PDCA1^-^F4/80^-^CD11c^+^CCR7^-^CD63^-^MHC-II^lo^ cells were then FACS-isolated into PBS+ 10% BCS on BD FACSAria^TM^ Fusion.

A single cell suspension of OT-II TCR transgenic (B6.Cg-Tg(TcraTcrb)425Cbn/J) thymocytes was prepared, and CD3^+^ CD4SP cells were FACS-isolated into PBS+10% BCS on the BD FACSAria^TM^ Fusion cell sorter. Thymocytes and DCs were seeded at a 2:1 ratio respectively, in a 96-well Round (U)-bottom plate in a total volume of 200µl complete RPMI per well. When indicated (Fig. 4g and Fig. 8c), 1µM OVA_323-339_ (GenScript) was included in the culture. 17.5µM CD40L blocking antibody (MR-1; BioXcell) or Isotype Control antibody (Catalog #: BE0091; BioXcell) were added to the indicated culture wells. Cultures were incubated for 16-18 hours at 37°C and then analyzed by flow cytometry to quantify activated DCs (CCR7^+^CD63^+^MHC-II^hi^ aDCs).

### Analysis of DC subsets in secondary lymphoid organs

LNs (combined inguinal, axillary, and brachial) and spleens from WT, *Rag2^-/-^*, *MHC-II^-/-^*, and *B2m^-/-^*mice were enzymatically digested with liberase as per the thymus protocol above. 6 x10^6^ cells were immunostained with the antibodies in Extended Data Fig. 2a for 40 minutes at 37°C prior to washing and resuspending in FWB+EDTA with 0.1μg/mL propidium iodide (Enzo). Analysis was performed on the BD FACSAria^TM^ Fusion.

### In vivo DC migration assay

To assay for WT versus *B2m^-/-^* DC migration into WT thymi, spleens from CD45.2^+^ WT and *B2m^-/-^* mice were enzymatically digested as above, and RBCs were lysed in RBC lysis buffer (Invitrogen). Splenocytes were enriched for DCs as described above. A 1:1 mixture of 5-10 x 10^6^ CD11c-enriched splenocytes from CD45.2^+^ WT and *B2m*^-/-^ mice each were resuspended in PBS prior to retro-orbital injection into non-irradiated hosts. CD45.1^+^ recipients were depleted for NK cells, 24 hours prior to DC transfer with αNK1.1 (PK136, BioXCell). Flow cytometry analysis was performed after 48 hours to identify transferred DCs. CD45.2 expression was used to distinguish donor from host thymic DCs, and MHC-I expression was used to distinguish WT from *B2m*^-/-^ donor DCs. 1-2 x 10^6^ cells from the 1:1 mixture above were immunostained for DC subsets on the day of the DC transfer to calculate the number of injected DCs within each subset and genotype. To calculate DC migration efficiency, the number of thymic DCs recovered following 48 hours within each subset and genotype was calculated relative to the number of injected DC subsets and genotypes, via flow cytometry.

To assay for the efficiency of WT DC migration into WT versus *B2m^-/-^*recipients, 5-10 x 10^6^ CD45.1^+^ splenic DCs were enriched, as above, and injected retro-orbitally into non-irradiated CD45.2^+^ WT or CD45.2^+^ *B2m^-/-^* recipients. An aliquot of enriched DCs were analyzed by flow cytometry to quantify the number of each DC subset transferred into recipient mice. After 48 hours, recipient thymi were enzymatically digested, as above, and cells were analyzed by flow cytometry to calculate the frequency of each donor DC subset recovered, relative to the input DCs. Donor and host DCs were distinguished based on CD45.2 versus CD45.1 expression, respectively.

### Generation of mixed and reciprocal BM chimeras

Long bones were extracted from F1, WT, *MHC-II^-/-^* and *B2m^-/-^* mice and were crushed in FWB using a mortar and pestle. Cells were passed through a 70µM cell strainer and immunostained with lineage-specific antibodies targeting CD11b, Gr1, Ter119, B220, CD3, CD4, CD8, prior to depletion with Dynabeads Sheep anti-rat IgG (Invitrogen). For mixed BM chimeras, 10 x 10^6^ lineage-depleted BM cells of WT and *MHC-II^-/-^* origin were injected at a 1:1 ratio into WT recipient mice that had been lethally irradiated (two split doses of 6Gy, separated by 3 hours). For reciprocal BM chimeras, 10 x 10^6^ BM cells of WT or *B2m*^-/-^ origin were injected into lethally irradiated WT or *B2m^-/-^* mice. In one reciprocal chimera experiment, NK cells were depleted, as described above, prior to BM transplantation; no significant differences from experiments without NK depletion were observed. Thymi from recipient mice were analyzed 10 weeks after reconstitution.

### qRT-PCR analysis of IFN gene expression by mTEC^hi^ cells

Thymi from WT and *B2m*^-/-^ mice were enzymatically digested as described above. and depleted for CD45^+^ cells with CD45-MicroBeads (Miltenyi Biotec) and LS columns as per the manufacturer’s protocol. CD45-depleted thymocytes were immunostained with antibodies against CD80, CD45, EpCAM, MHC-II, CD11c, Ly51, and with the lectin UEA-1 (Vector Laboratories). 1-3 x10^4^ mTEC^hi^ (CD80^+^EpCAM^+^UEA-1^+^Ly51^-^MHC-II^hi^) cells were FACS-sorted directly into Trizol (Invitrogen) and stored at -80°C.

cDNA conversion was performed with the SuperScript IV VILO kit (Invitrogen) according to manufacturer’s instructions. SYBR green (Applied Biosystems) was used to quantify product amplification and samples were analyzed on the Quant7 Flex RT-PCR system (ThermoFisher). Samples were run in technical triplicates and averaged ΔΔCt values were calculated relative to beta-actin expression within each sample and by comparison to paired WT mTEC^hi^ samples. Primers used are as follows: *beta-actin* (forward: CACTGTCGAGTCGCGTCCA, reverse: CATCCATGGCGAACTGGTGG). : *Ifnb1*(forward: CCCTATGGAGATGACGGAGA, reverse: CTGTCTGCTGGTGGAGTTCA), *Ifnl2* (forward: AGGTGCAGTTCCCACCTCT, reverse: TCAGTCATGTTCTCCCAGACC), *Ifnl3* (forward: TCCCAGCTGCGACCTGT, reverse: CAGGGGTCTCCTTGCTCTG).

### Statistical analyses

All statistical tests were carried out using Prism (V9.2.0; GraphPad Software) and significance was determined using 1-way analysis of variance (ANOVA) with Tukey’s multiple comparisons test/Dunnett’s multiple comparisons test, or unpaired Student *t*-test, as indicated in the figure legends. p < 0.05 was considered significant: *p < 0.05; **p < 0.01; ***p < 0.001; ****p < 0.0001. Number of replicates and sample size for each experiment are included in the corresponding Figure legends. All statistical tests for histocytometry analysis were performed with R 4.3.1.

## DATA AND CODE AVAILABILITY

scRNA-seq data have been deposited at the Gene Expression Omnibus (GEO: GSE239780). This study generated original code for histocytometry analyses, which can be found via Github (https://github.com/EhrlichLab/histocytometry_Srinivasan_2023). Scripts used for p-Creode analyses are available via Github (github.com/Ken-Lau-Lab/pCreode). Any additional information required to reanalyze the data reported in this paper is available upon request from the lead contact, Lauren I.R. Ehrlich.

## Supporting information

Supplemental Data Srinivasan et al.

## ACKNOWLEDGEMENTS

We thank the Animal Resource Center staff at the University of Texas at Austin for mouse maintenance support and Richard Salinas for maintenance of FACS instruments. scRNA-seq library preparation and sequencing was performed by Holly Stevenson at the Genomic Sequencing and Analysis Facility at UT Austin, Center for Biomedical Research Support. *Xcr1^cre/+^Cd40^f/-^* and *Cd40^-/-^* mice were generously provided by Dr. Kenneth Murphy. We thank Drs. Nancy Manley, Ellen Richie, and. Laura Hale for constructive feedback and Dr. Stephen Yi for initial bioinformatics analyses. This research was supported by a grant from the National Institutes of Health, P01AI139449, to L.I.R.E., K.L., and Q.L. Servers contributed by Advanced Micro Devices (AMD) to L.I.R.E. were used for imaging data analysis.

## AUTHOR CONTRIBUTIONS

L.I.R.E, J.S., C.M., and A.C. conceptualized the study and wrote the manuscript. J.S., C.M., and A.C. designed and performed experiments and analyzed data with supervision from L.I.R.E. K.L., Q.L., B.H., Y.Y., and C.H. analyzed scRNA-seq datasets. J.M. carried out histocytometry analyses. H.J.S. assisted with FACS-isolation of cells for functional assays. L.I.R.E. acquired funding.

