## Supplemental Data Srinivasan et al. for "Single-cell transcriptomics reveals heterogenous thymic dendritic cell subsets with distinct functions and requirements for thymocyte-regulated crosstalk"

### Extended Data Figure 1

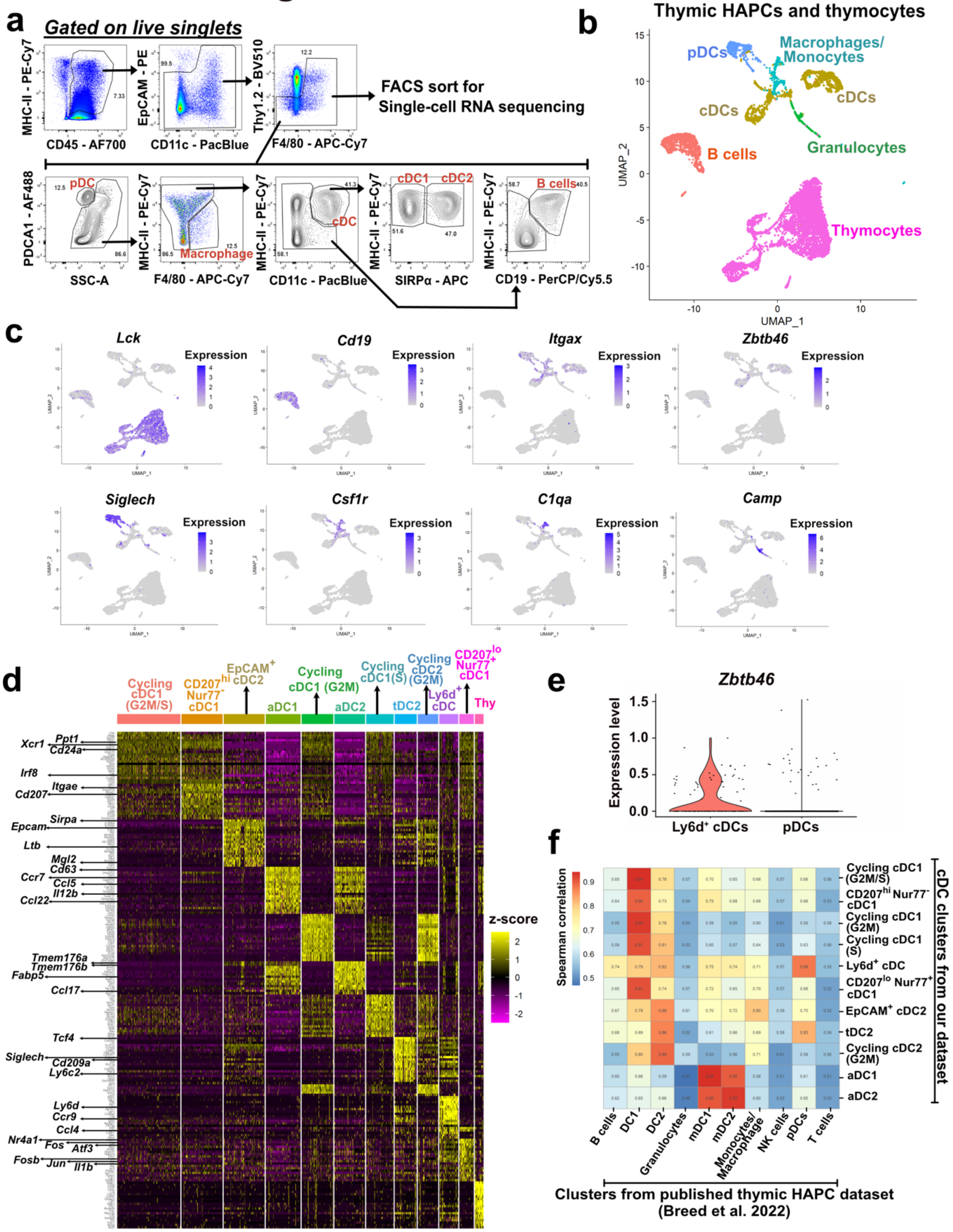

**Extended Data Fig. 1. Identification of transcriptionally distinct HAPC subsets, related to Fig. 1.**

(a) Flow cytometric gating strategy used to sort HAPCs for scRNA-seq analysis. (b) UMAP visualization showing major HAPC populations identified from 8404 FACS-sorted HAPCs (and residual thymocytes) from adult C57BL6/J thymi- B cells (*Cd19*), cDCs (*Zbtb46, Itgax*), pDCs (*Itgax, Siglech*), macrophages/monocytes (*Csf1r, C1qa*), granulocytes (*Camp*), and thymocytes (*Lck*). (c) Feature plots showing expression levels of select genes associated with the major HAPC subsets overlaid on the UMAP visualization from (b). (d) Heatmap displaying normalized expression of the top 10 differentially expressed genes in each transcriptionally distinct cDC cluster from Fig. 1a. Arrows highlight select genes specific to the transcriptionally distinct cDC1, cDC2, and aDC clusters. (e) Violin plot displaying normalized expression levels of *Zbtb46* gene in Ly6d$^{+}$ cDCs and pDCs. (f) Spearman correlation analysis of thymic cDC clusters from our dataset, and cDC and other clusters in the dataset from [11].

### Extended Data Figure 2

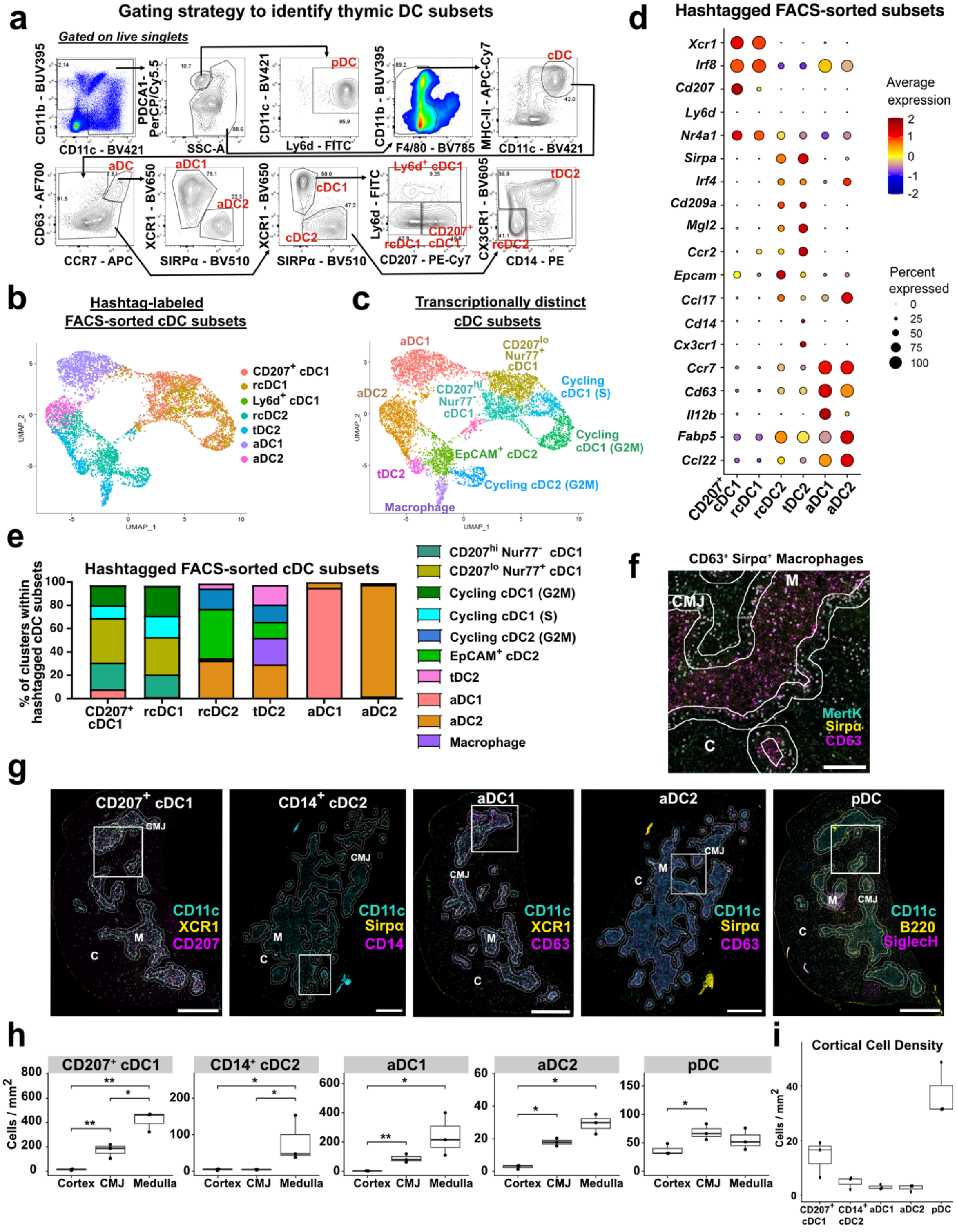

**Extended Data Fig. 2. Identification of transcriptionally distinct cDC clusters and their correlation with FACS-sorted cDC subsets, related to Fig. 1.**

(a) Flow cytometric gating strategy to identify distinct DC subsets shown in Fig. 1d. (b) UMAP visualization of scRNA-seq transcriptional data from pooled FACS-sorted cDC subsets, colored by their hashtag labels. (c) Same UMAP plot as (b), colored by transcriptionally distinct cDC clusters defined by the Seurat pipeline. (d) Dot plot showing normalized expression levels (z-score) of select genes associated with the hashtagged FACS-sorted cDC subsets. Expression is normalized per row (representing a gene) across different FACS-sorted cDC subsets. (e) Frequency of each transcriptionally distinct cDC cluster contained within the FACS-sorted hashtagged cDC subsets. (f) Representative immunofluorescence image of $MertK^{+}$ macrophages and $CD63^{+}SIRP\alpha^{+}MerTK^{-}$ aDC2s in thymic sections (representative of N=3 independent experiments). Centroids are placed over $CD63^{+}$ $SIRP\alpha^{+}$ cells that also express MertK: the frequency of MertK-expressing cells among $CD63^{+}SIRP\alpha^{+}$ myeloid cells was used to correct aDC2 densities from data as in (h). Scale bar, 250µm. Lines demarcate the cortex (C), medulla (M), and CMJ in the thymic sections. Centroids with 40% opacity were overlaid on the quantified cells. (g) Representative immunofluorescence images of DC subsets (representative of N=3 independent experiments), identified by the indicated markers. White boxes indicate the regions of interest displayed in Fig. 1e. Scale bars, 1,000µm. (h) Quantification of cell densities of each of the indicated DC subsets in distinct anatomical regions of the thymus, based on histocytometry analyses of data as in (f-g). Data are compiled from N=3 independent experiments. Box plots represent mean ± SD. Statistical analysis was performed using Tukey's Honest Significant Difference test, where $*p<0.05$, $**p<0.01$. (i) Quantification of cortical cell densities of each of the indicated DC subsets, based on histocytometry analyses of data as in (f-g). Data are compiled from N=3 independent experiments. Box plots represent mean ± SD.

### Extended Data Figure 3

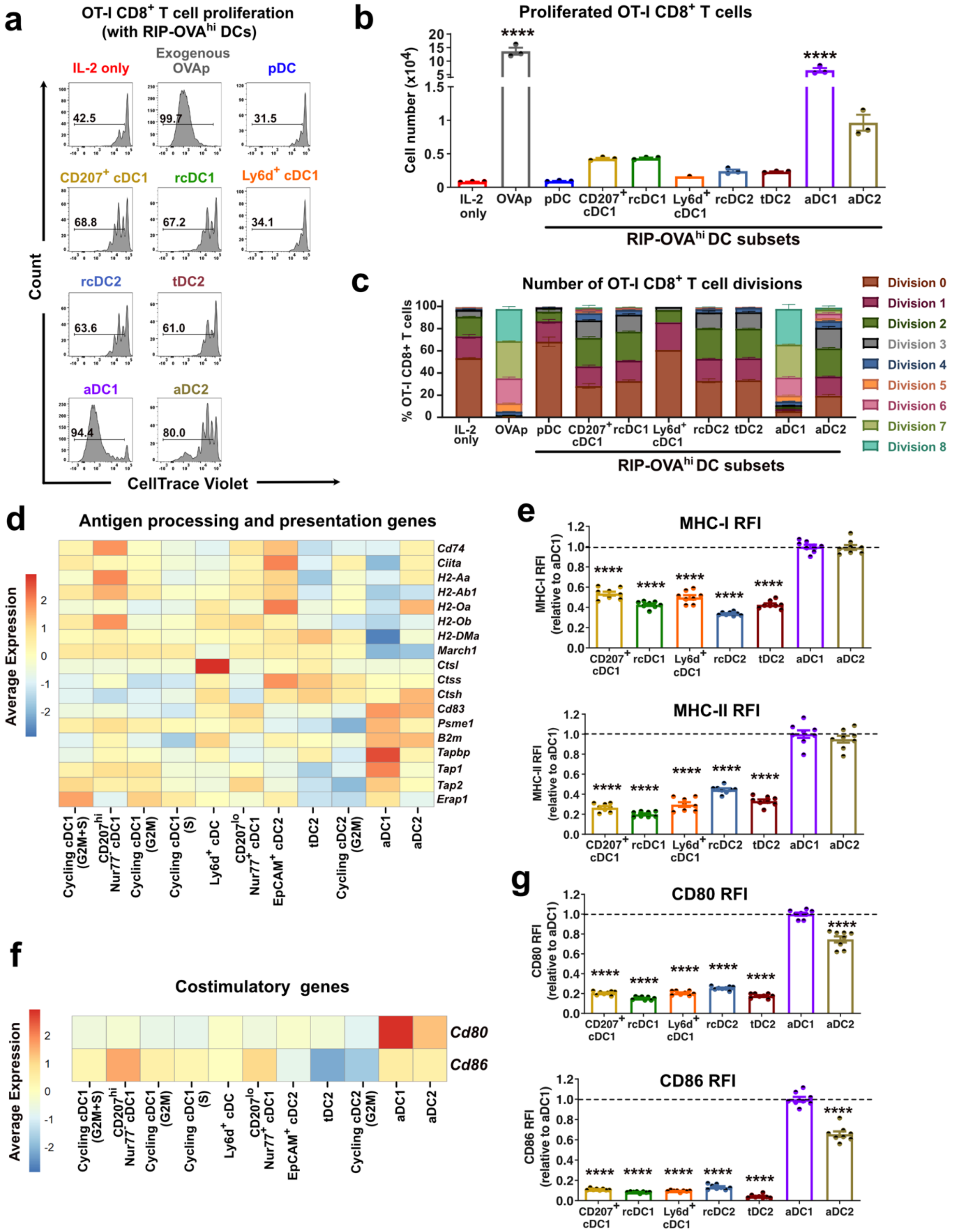

**Extended Data Fig. 3. aDC1s, followed by aDC2s, are highly efficient at mTEC-derived antigen presentation on MHC-I and MHC-II, related to Fig. 2.**

(a) Representative histograms showing the percentage of OT-I $CD8^+$ T cells that underwent at least 1 cell division after co-culture for 3.5 days with the specified DC subsets isolated from RIP-$OVA^{hi}$ mice. OT-I $CD8^+$ T cells incubated only with IL-2 served as negative controls, while those co-cultured with OVA-pulsed splenocytes (50nM $OVAp_{257-264}$) served as positive controls. For (a-c), data are representative of 3 independent experiments. (b) Quantification of OT-I $CD8^+$ T cells that underwent proliferation from the experiment described in (a). Bars represent mean ± SEM. Each dot represents a replicate well. Statistical analysis was performed using one-way ANOVA with Dunnett's multiple comparisons test, where ****p<0.0001. Significance is relative to "IL-2 only" control wells. (c) Modeling, based on data from the experiment in (a), of the frequency of OT-I $CD8^+$ T cells that underwent the indicated number of cell divisions. Bars represent mean ± SEM. (d, f) Heatmap displaying average normalized expression (z-score) of select (d) MHC-I and MHC-II antigen processing and presentation genes and (f) costimulatory genes in each cDC cluster from Fig. 1a. Expression is normalized per gene across different cDC clusters. (e, g) Relative cell-surface expression levels of (e) MHC-I and MHC-II, and (g) CD80 and CD86 proteins by cDC subsets, as determined by flow cytometry. Data are normalized to average MFI levels of aDC1s for each protein within an individual experiment. Data are compiled from N=3 independent experiments (n=8 mice). Bars represent mean ± SEM and symbols represent individual mice. Statistical analysis was performed using one-way ANOVA with Dunnett's multiple comparisons test, where ****p<0.0001. Significance is relative to the aDC1 subset.

### Extended Data Figure 4

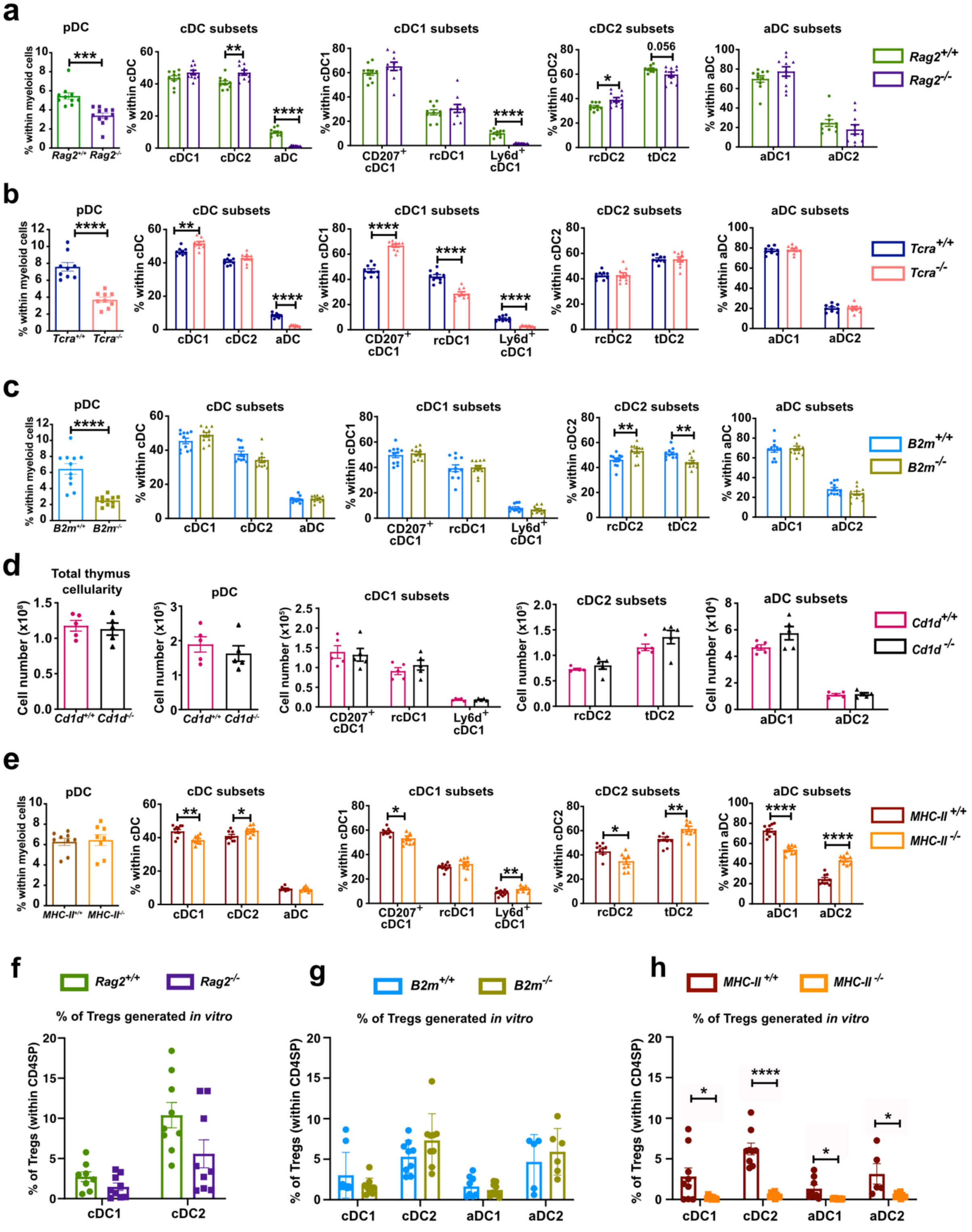

**Extended Data Fig. 4. Thymic cDC1 and cDC2 homeostasis and activation depend on the presence of distinct thymocyte subsets, related to Fig. 3.**

(a-c) Quantification of (left to right) frequencies of pDCs, total cDC subsets, cDC1 subsets, cDC2 subsets, and aDC subsets in the thymi of (a) *Rag2*$^{-/-}$, (b) *Tcra*$^{-/-}$, and (c) *B2m*$^{-/-}$ relative to littermate control mice. The population from which the frequencies were calculated is indicated on the respective Y-axes. (d) Quantification of (left to right) total thymus cellularity and cell numbers of pDCs, cDC1 subsets, cDC2 subsets, and aDC subsets of WT versus *Cd1d*$^{-/-}$ littermate mice. (e) Quantification of (left to right) frequencies of pDCs, total cDC subsets, cDC1 subsets, cDC2 subsets, and aDC subsets in the thymi of *MHC-II*$^{-/-}$ relative to littermate control mice. (a-e) Data are compiled from N=3-4 independent experiments (n=8-11 mice). Bars represent mean ± SEM and symbols represent individual mice. (f-h) Quantification of frequencies of Foxp3$^{+}$ CD25$^{+}$ Tregs induced after co-culturing CD73$^{-}$ CD25$^{-}$ Foxp3$^{-}$ CD4SP thymocytes with the indicated thymic DC subsets from WT versus (e) *Rag2*$^{-/-}$, (f) *B2m*$^{-/-}$, and (g) *MHC-II*$^{-/-}$ mice for 4 days. Data are compiled from N= 3 independent experiments. Bars represent mean ± SEM and symbols represent individual wells per DC subset from N=3 experimental repeats. Statistical analysis was performed using unpaired Student *t*-test; *p<0.05, ** p<0.01, ***p<0.001, ****p<0.0001.

### Extended Data Figure 5

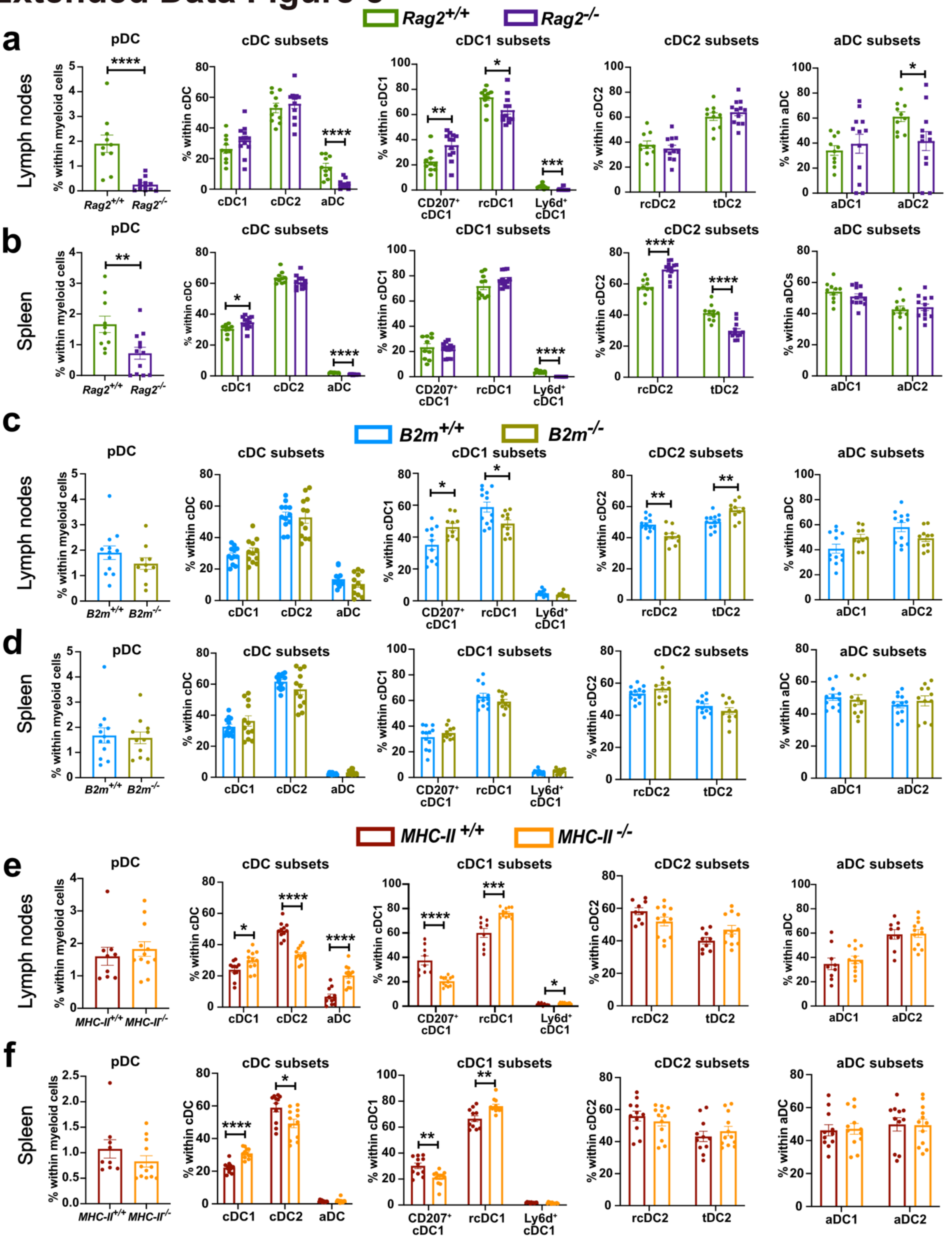

**Extended Data Fig. 5. Quantification of DC subsets from lymph nodes and spleens of mice with impaired T cell development, related to Fig. 3**

(a-f) Quantification of frequencies of (left to right) pDCs, cDC subsets, cDC1 subsets, cDC2 subsets and aDC subsets in (a,c,e) pooled axillary, brachial, and inguinal lymph nodes and (b,d,e) spleens of (a-b) *Rag2*$^{-/-}$ , (c-d) *B2m*$^{-/-}$, and (e-f) *MHC-II*$^{-/-}$ mice relative to WT controls. The populations from which the frequencies were calculated are indicated on the Y-axes. (a-f) Data are compiled from N=3 independent experiments (n=9-12 mice). Bars represent mean ± SEM and symbols represent individual mice. Statistical tests were performed using unpaired Student t-test; *$p<0.05$, ** $p<0.01$, ***$p<0.001$, ****$p<0.0001$.

### Extended Data Figure 6

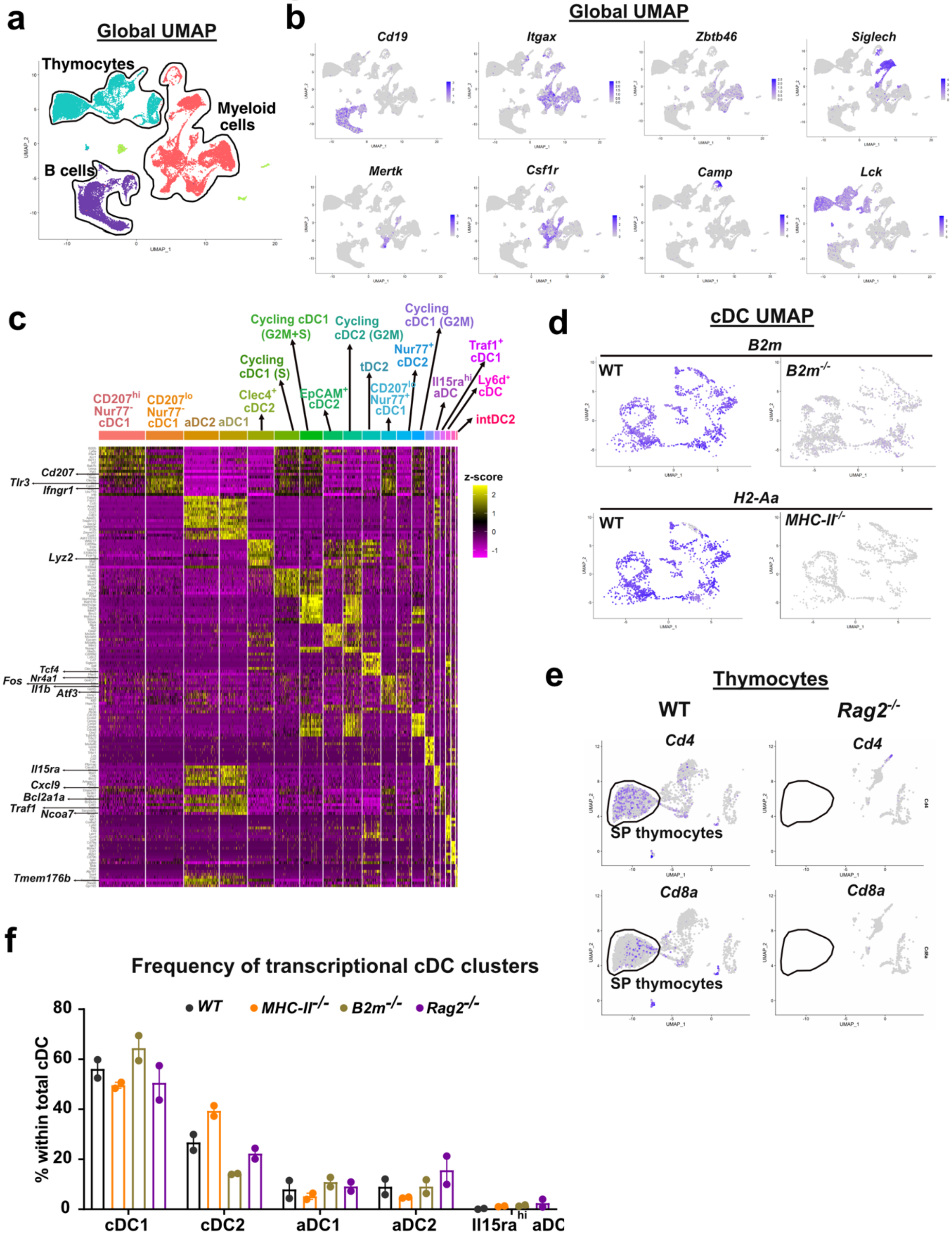

**Extended Data Fig. 6. Transcriptional profiling of thymic HAPCs in mice devoid of distinct thymocyte subsets reveals the impact on cellular transcriptomes and frequencies, related to Fig. 5.**

(a) Merged UMAP visualization of 34,750 cells depicting major cell continents in WT, *MHC-II*$^{-/-}$, *B2m*$^{-/-}$ and *Rag2*$^{-/-}$ HAPC scRNA-seq datasets. Data are compiled from N=2 independent experiments (n=2 mice). (b) Feature plots displaying expression levels of select genes associated with distinct HAPC subtypes – B cells (*Cd19*), cDC (*Itgax, zbtb46*), pDC (*Siglech, Itgax*), Macrophages and Monocytes (*MertK, Csf1r*) and granulocytes (*Camp*). Residual thymocytes included in the sorted cells were identified by *Lck* expression. (c) Heatmap displaying normalized expression of the top 10 enriched genes in each transcriptionally distinct cDC cluster from Fig. 5a. (d) Feature plots showing scRNA-seq expression levels of (top) *B2m* transcripts in WT versus *B2m*$^{-/-}$ cDCs and (bottom) *H2-Aa* transcripts in WT versus *MHC-II*$^{-/-}$ cDCs overlaid on the UMAP visualization from Fig. 4a. (e) Feature plots showing expression levels of *Cd4* and *Cd8a* transcripts in thymocytes from WT versus *Rag2*$^{-/-}$ mice overlaid on the UMAP visualization in (a). (f) Quantification of the frequencies of cDC1, cDC2, aDC1, aDC2, and Il15ra$^{hi}$ aDC subsets identified in Fig. 4a from scRNA-seq transcriptional profiling of thymic DCs in the four indicated genotypes. Data are compiled from 2 independent experiments per genotype (n=2 mice). Bars represent mean ± SEM and symbols represent individual mice.

#### Extended Data Figure 7

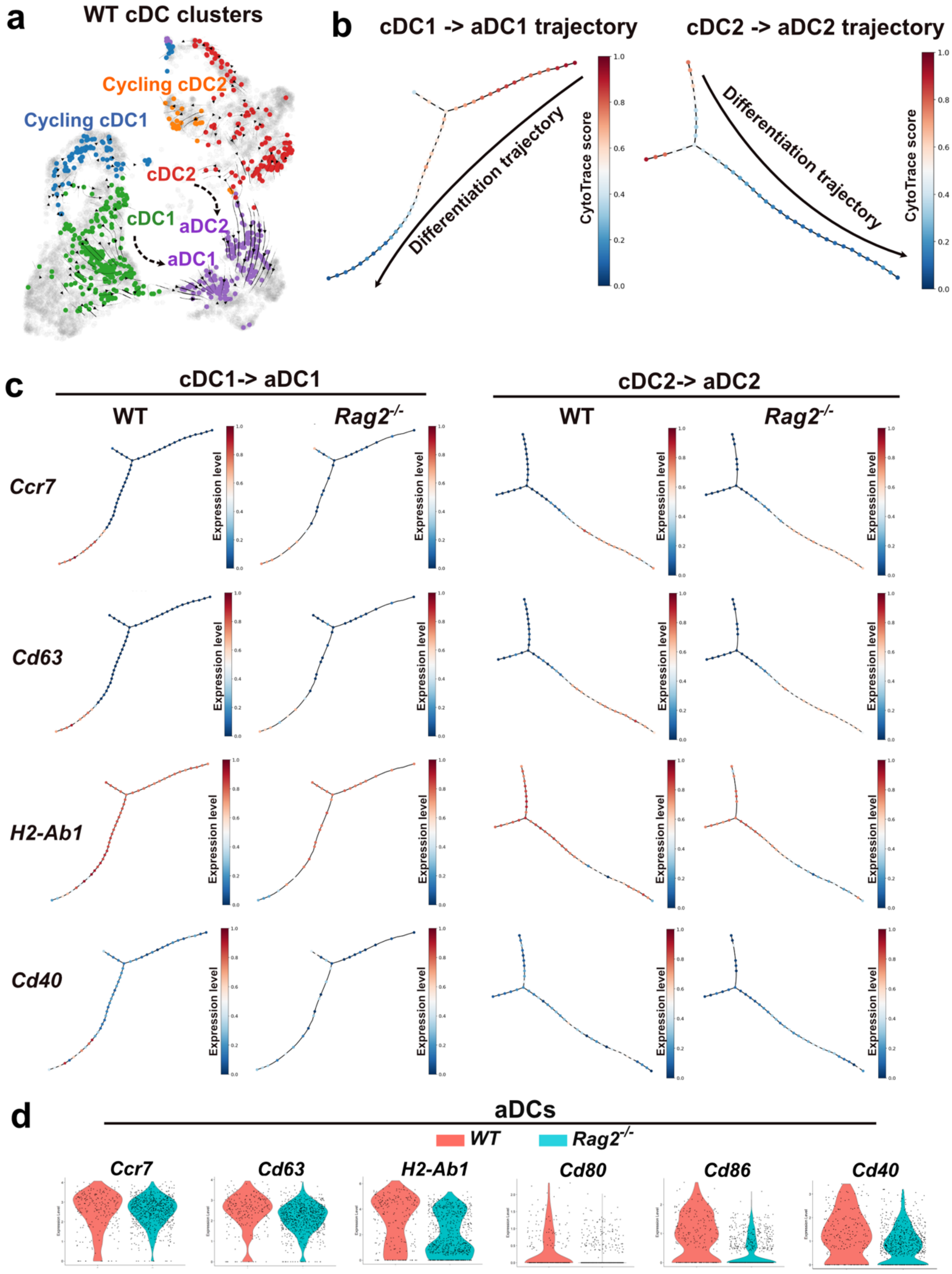

**Extended Data Fig. 7. SP thymocytes are necessary for full cDC activation, related to Fig. 6.**

(a) RNA velocity analysis with arrows showing relative transition probabilities derived from ratio calculations of unspliced over spliced transcripts (cDCs -> aDCs). (b) p-Creode analysis depicting cell-state transition trajectories for (left) cDC1 and (right) cDC2 transcriptional subsets. The topology was constructed with hierarchical placement, which connects the two transcriptionally closest end-states with path nodes. Overlay represents CytoTrace score to indicate cells with a high degree of stemness and transcriptional diversity (score closer to 1) versus those with a low degree of stemness, indicative of a more differentiated transcriptional state (score closer to 0). (c) p-Creode (left) cDC1 and (right) cDC2 differentiation trajectories from WT and *Rag2*$^{-/-}$ scRNA-seq datasets overlaid with normalized expression values of select genes associated with cDC activation and licensing. (d) Violin plots showing normalized expression levels of the indicated genes, associated with DC activation or costimulation of T cells, in WT versus *Rag2*$^{-/-}$ aDCs (combined aDC1, aDC2, and Il15ra$^{hi}$ aDC clusters).

### Extended Data Figure 8

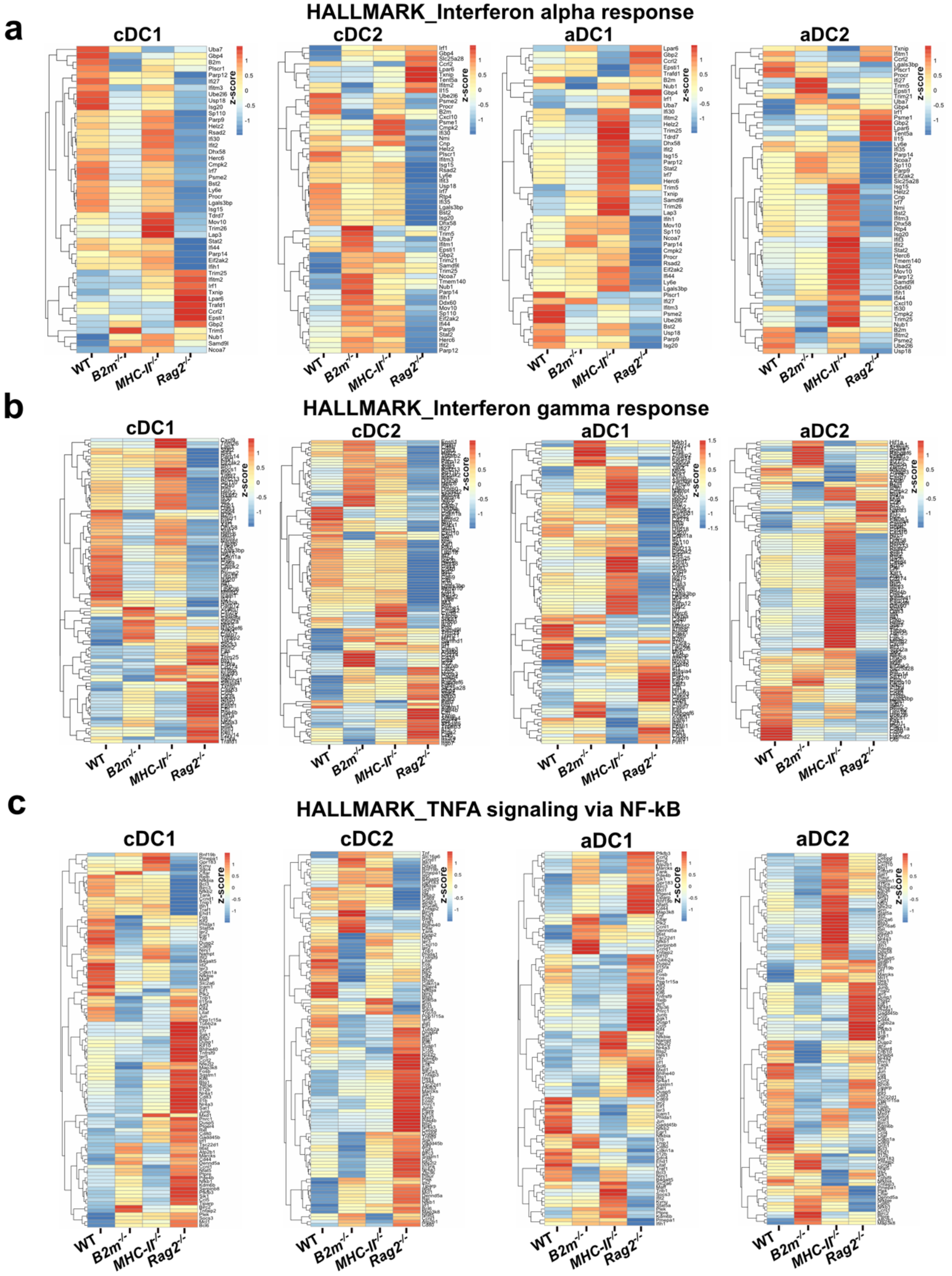

**Extended Data Fig. 8. CD8SP thymocytes promote expression of genes associated with interferon signaling in DCs, while DCs activated in the absence of SP thymocytes express elevated levels of genes associated with canonical NF-kB signaling, related to Fig. 6.**

(a-c) Heatmaps show normalized expression (z-score) of genes associated with the indicated pathways in cDC1s, cDC2s, aDC1s and aDC2s. Genes shown represent the union of differentially regulated genes identified via tradeSeq analysis along the p-Creode cDC1 and cDC2 differentiation trajectories from pairwise WT versus KO comparisons (Fig. 6c-d). Pathways were identified via gene enrichment analysis of the differentially regulated genes along the cDC1/cDC2 trajectories, carried out using WebgestaltR with pathways from KEGG and MSigDB databases.

### Extended Data Figure 9

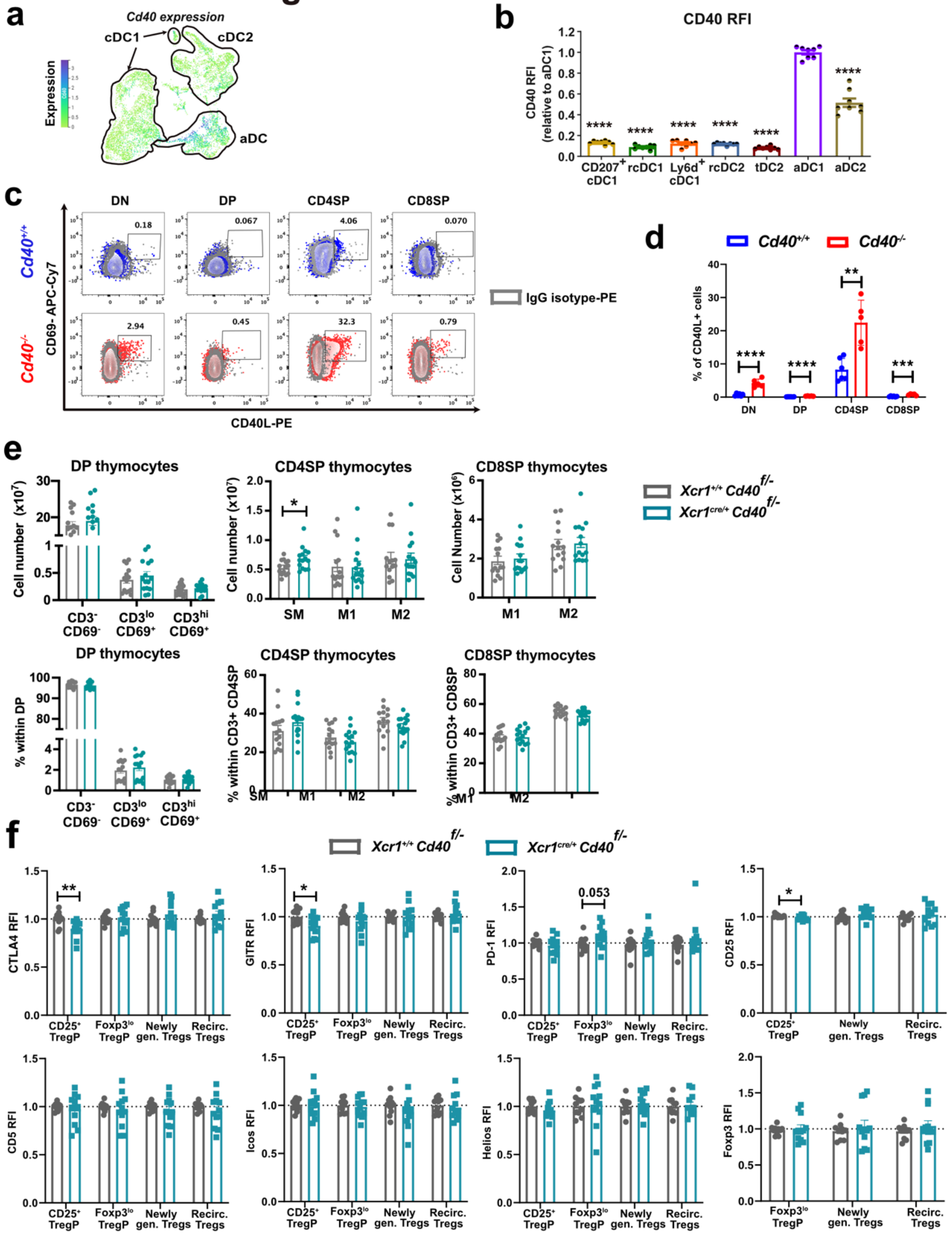

**Extended Data Fig. 9. Cognate interactions with CD4SP thymocytes and CD40-CD40L interactions are indispensable for cDC1 activation and central tolerance, related to Fig. 8**

(a) *Cd40* gene expression levels overlaid on the UMAP representation of scRNA-seq datasets from WT and KO thymic cDCs, as in Fig. 5a. CD40 expression is shown on cDC and aDC subsets from the KO DC UMAP. (b) CD40 RFI levels in flow cytometry-defined cDC subsets. Data are normalized to average MFI levels of aDC1s per experimental repeat. Data are compiled from N=3 independent experiments (n=8 mice) (c-d) (c) Representative flow cytometry plots and (d) quantification of the frequencies of CD40L expression by the indicated major thymocyte subsets. Bulk thymocyte populations are plotted from the same experiments as for Fig. 8b with N=2 independent repeats (n=5-6). (e) Quantification of cell numbers (top) and frequencies (bottom) of the indicated (left to right) DP, CD4SP, and CD8SP thymocyte subsets in $Xcr1^{+/+}CD40^{f/-}$ versus $Xcr1^{cre/+}CD40^{f/-}$ littermate mice. Data are compiled from N=5 independent experiments (n=14-15 mice). (f) The expression of Treg effector markers in $CD25^{-}Foxp3^{-}$ CD4SP conventional thymocytes, $CD25^{+}Foxp3^{-}$ TregP, $CD25^{-}Foxp3^{lo}$ TregP, and $CD25^{+}Foxp3^{+}$ Tregs in $Xcr1^{+/+}CD40^{f/-}$ versus $Xcr1^{cre/+}CD40^{f/-}$ thymi. Data are compiled from N=4 independent experiments (n=11-12 mice). (b,d,e,f) Bars represent mean ± SEM and symbols represent individual mice. Statistical tests were performed using (b) one-way ANOVA with Dunnett's multiple comparisons test relative to aDC1 and (d-g) unpaired Student t-test. *p<0.05, ** p<0.01, ***p<0.001, ****p<0.0001.
